# A hierarchy of causes of death in senescent *C. elegans*

**DOI:** 10.1101/2025.08.21.671442

**Authors:** Hongyuan Wang, Carina C. Kern, Chiminh Nguyen Hong, Alis Saez Allende, Jiayi Qiao, Aihan Zhang, Yimu Fan, Marina Ezcurra, David Gems

**Affiliations:** Institute of Healthy Ageing, and Research Department of Genetics, Evolution and Environment, University College London, London WC1E 6BT, UK; School of Biosciences, Stacey Building, University of Kent, Canterbury, UK

**Keywords:** Aging, *C. elegans*, Competing risks, Infection, Mortality, Pathology, Teratoma, Tumor

## Abstract

In experimental gerontology, lifespan is often interpreted as a metric of the rate of the overall aging process. However, interventions that increase lifespan can result from suppression of one or more individual late-life pathologies. Here we show how, in the nematode *Caenorhabditis elegans*, such pathologies can compete in a hierarchical fashion to cause death, such that removal of one cause of death can unmask another. Under standard culture conditions, a major cause of death in elderly *C. elegans* is infection by their bacterial food source. We report that only when such infection is prevented is lifespan extended by suppression of a second senescent pathology, teratoma-like uterine tumors. Thus, as in mammals, lifespan in wild-type *C. elegans* can be limited by naturally-occurring neoplasia. By contrast, blocking bacterial infection attenuated the life-shortening effects of vitellogenesis, and did not unmask a life-shortening effect of distal gonad degeneration. Thus, depending on the masking or unmasking of competing causes of mortality in the hierarchy of causes of death, nematode lifespan limitation in different contexts can reflect action of distinct life-limiting senescent pathologies. This underscores how increases in lifespan do not necessarily reflect a reduction in overall aging rate.

## Introduction

Although there is little consensus within the field of biogerontology about the causes of senescence (aging), it is at least clear that its principal, ultimate cause is the process of evolution (Rose, 1991; Williams, 1957). Evolutionary physiology seeks to understand the evolved proximate mechanisms by which genes cause senescence (Arnold and Rose, 2023; Gems and Kern, 2024). Research guided by an emerging explanatory framework, programmatic theory, has yielded useful insights into the causes of senescence in the short-lived nematode *C. elegans* (de la Guardia et al., 2016; Ezcurra et al., 2018; Galimov and Gems, 2020; Gems and de la Guardia, 2013; Gems et al., 2021; Kern et al., 2023; Kern et al., 2021; Slade et al., 2024; Wang et al., 2018b).

According to one concept within programmatic theory, lifespan is limited by hyperfunctional (i.e. excessive) levels of wild-type mechanistic target of rapamycin (mTOR) signaling (Blagosklonny, 2006). The originator of the hyperfunction theory, Misha Blagosklonny, suggested that although stochastic molecular damage accumulates throughout life, this is not a major, primary cause of late-life mortality. This is because mTOR hyperfunction causes death before damage accumulation can substantially promote life-limiting senescent pathology (with the exception of cancer) (Blagosklonny, 2008). Thus, the hyperfunction theory includes a competing risks view of the causes of lifespan limitation by senescence.

The evolutionary biologist George C. Williams once asked the senior author of the present study: “What do senescent *C. elegans* die from?” While aging people die mainly from cardiovascular disease, cancer and chronic obstructive pulmonary disease, the causes of death in aging *C. elegans* are less well understood. However, it is clear that death for elderly *C. elegans* under standard culture conditions (Brenner, 1974) is a consequence of a combination of extrinsic and intrinsic factors.

A major extrinsic cause is infection with the *Escherichia coli* bacteria that is supplied as a food source, to which all individuals succumb in later life (Garigan et al., 2002; Gems and Riddle, 2000; Podshivalova et al., 2017). This occurs due to a combination of at least two intrinsic changes: mechanical senescence of the pharyngeal cuticle in early adulthood, and later processes of immune senescence (Zhao et al., 2017). Another possible internal cause is vitellogenesis (yolk synthesis), whose inhibition by RNA-mediated interference (RNAi) reduces several senescent changes, including intestinal atrophy and formation of pseudocoelomic lipoprotein pool (accumulations of lipid and yolk in the body cavity), and also extends lifespan (Ezcurra et al., 2018; Murphy et al., 2003; Sornda et al., 2019). However inhibition of several other major senescent pathologies does not increase lifespan under standard culture conditions: degeneration and fragmentation of the distal gonad (de la Guardia et al., 2016), and formation of uterine tumors (Riesen et al., 2014).

In this study, we explore a competing risks hypothesis of *C. elegans* lifespan determination (Figure 1A, left), in which causes of mortality are, to some degree, nested, in a manner akin to a Russian matryoshka doll, or the nested skins of an onion. According to this hypothesis, extrinsic causes contribute particularly to the outer skin, underneath which lie partially or fully masked intrinsic causes. Arguably, first among the latter will come programmatic causes of mortality (e.g. due to hyperfunction). Then, ultimately, according to Blagosklonny’s scheme, after programmatic causes will come accumulation of stochastic molecular damage. According to this model, the identity of the senescent pathologies that limit lifespan is context dependent (Ezcurra et al., 2018). Here a counter-hypothesis, and a traditional view (at least, implicitly), is that senescence is, as it were, all one thing, happening all at once, and limiting life (Figure 1A, right).

**Figure 1.**
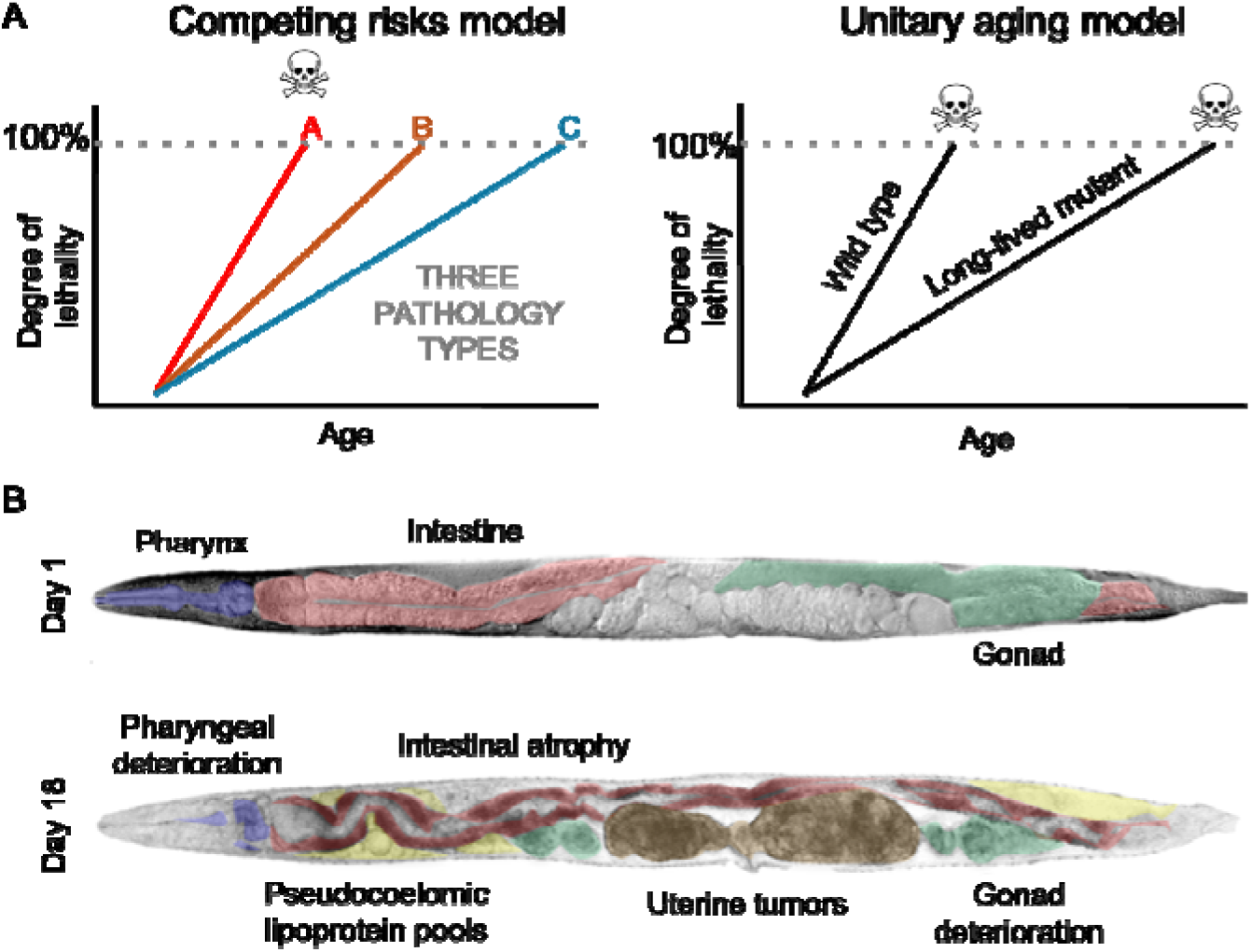
(A) Left, the competing risks model (standard culture conditions). Senescent pathologies develop, gradually become more severe, until they become lethal, causing death. Here pathology A is life limiting. If A is prevented, then pathology B becomes life limiting. Right, A counter-hypothesis, where a unified overall process of senescence limits life. Only according to the right hand scheme can lifespan be viewed as a metric of the overall aging process. (B) Senescent pathologies in *C. elegans* hermaphrodites, including uterine tumors (brown). Top, young adult hermaphrodite, with no uterine tumors (day 1 of adulthood). Bottom: Elderly hermaphrodite showing typical paired uterine tumors, one in each branch of the uterus. Several other pathologies are also visible.

Among the most striking pathologies in senescent *C. elegans* hermaphrodites are the uterine tumors (Figure 1B, bottom), that frequently grow so large as to fill the entire width of the body cavity in the mid-body region. Uterine tumors (also referred to as oocyte clusters) develop in all hermaphrodites after depletion of self-sperm stocks (Luo et al., 2010). Completion of meiosis II in terminal oocytes requires fertilization, and in its absence, diploid oocytes are ovulated into the uterus. There they undergo numerous rounds of DNA endoreduplication, leading to formation of disordered chromatin masses, and to cellular hypertrophy (Golden et al., 2007).

Older uterine tumors show signs of recapitulation of later embryonic gene expression (Wang et al., 2018b), suggesting that their formation represents a futile attempt by diploid oocytes to execute a program of embryogenesis (or embryogenetic quasi-program). This and the similarity in etiology to that of ovarian teratomas (ovarian cysts, arising from failure of meiosis II) in female mammals suggests that uterine tumors are a form of teratoma (Wang et al., 2018a; Wang et al., 2018b).

Uterine tumor development can be prevented using 5-fluoro-2’-deoxyuridine (FUDR, or floxuridine), a drug used for the treatment of colorectal cancer. Surprisingly, under otherwise standard culture conditions, this does not increase lifespan, i.e. these large tumors have no effect on late-life mortality (Riesen et al., 2014). However, results of a recent study employing machine learning to assess the role of individual senescent pathologies in lifespan limitation suggest that under conditions that extend lifespan, uterine tumors may contribute to late-life mortality (Kern et al., 2025). Here we explore the competing risks model by examining effects of preventing senescent pathology development on late-life mortality under different conditions. Our results provide an example of a programmatic pathology, uterine tumor development, that contributes to late-life mortality in a context-dependent fashion.

## Results

### FUDR extends lifespan when bacterial infection is prevented

Human populations with higher death rates throughout life from infectious pathogens will experience concomitantly lower death rates from cancer (which predominantly occurs in later life). We wondered whether, by the same token, uterine tumors contribute to late-life mortality in the absence of bacterial infection. To this end, we first compared the effects on lifespan of suppressing tumor growth, using 50 μM FUDR, administered from the late-larval L4 stage or D2 (day 2) of adulthood, in the presence and absence of an antibiotic (carbenicillin, Carb). The following account describes summed data from three trials, unless otherwise stated; for full statistics, see Supplemental Tables; for raw data for all lifespan trials in this study, see Dataset S1).

In the absence of Carb, FUDR did not increase longevity: it slightly shortened mean lifespan when administered from L4 but not from D2 (respectively -6.1%, +6.0%, *p* = 0.0024, *p* = 0.27, log rank test; Figure 2A; Table S1). Antibiotic treatment extended lifespan (+52.2%, *p* < 0.0001, Figure 2A), as previously seen (Garigan et al., 2002). In the presence of Carb, FUDR from L4 again slightly shortened lifespan (-5.8%, *p* < 0.0001), but FUDR from D2 significantly increased it (+22.0%, *p* < 0.0001, Figure 2A). The life-shortening effects of FUDR from L4 may reflect a deleterious effect of drug on later development at this relatively high drug concentration. In all subsequent trials, FUDR was administered from D2 onwards unless otherwise stated.

**Figure 2.**
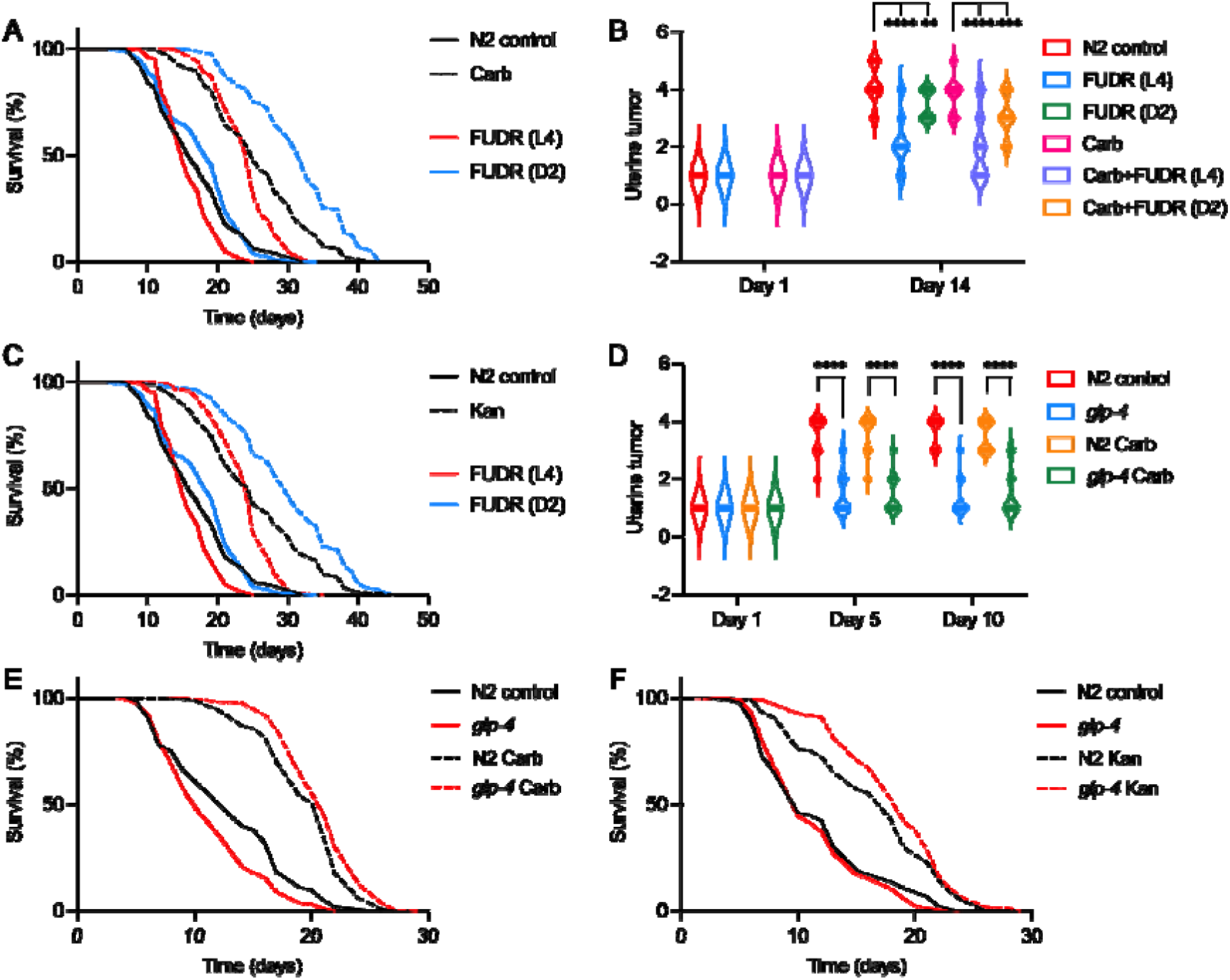
50 μM FUDR increases lifespan when bacterial infection is prevented. (A) FUDR from day 2 of adulthood (D2) increases lifespan in the presence of Carb. (B) FUDR suppresses uterine tumor development. (C) FUDR increases lifespan in the presence of Kan. (D) *glp-4(bn2)* suppresses uterine tumor development, with or without Carb. (E,F) *glp-4(bn2)* extends lifespan when bacterial infection is prevented using Carb (E), or Kan (F). For individual trials and statistical comparisons in lifespan analyses, see Table S1 (A, C), and Table S2 (E, F).

The life-extending effect of FUDR could be attributable to tumor suppression, amelioration of other senescent pathologies, or some other effect. To explore the second possibility, effects of FUDR on several major senescent pathologies were assessed: uterine tumor development, pharyngeal deterioration, distal gonad degeneration, and intestinal atrophy. In the presence of Carb, FUDR from D2 significantly reduced tumor development, but not the other pathologies (Figure 2B, Figure S1A, B).

To verify that the putative unmasking effect of Carb is attributable its antibiotic properties, we performed similar tests using an alternative means to block bacterial infection: a different class of antibiotic (kanamycin, Kan) which, like Carb, increases *C. elegans* lifespan (Garigan et al., 2002). Here again, in the presence of Kan, FUDR extended lifespan (+17.8%, *p* < 0.0001; Figure 2C, Table S1). These findings are consistent with prevention of mortality from bacterial infection unmasking effects of uterine tumors on late-life mortality. However, it remains possible that the life-extending effect of FUDR is caused by something other than tumor suppression.

### *glp-4(bn2)* extends lifespan when bacterial infection is prevented

One way to investigate this is to test whether suppression of tumor development by other means also increases lifespan when bacterial infection is prevented. *glp-4(bn2)* (<u>g</u>ermline <u>p</u>roliferation defective) is a temperature-sensitive mutation that blocks germline proliferation at 25°C but not 15°C. If raised from egg at 25°C, inhibition of germline signaling leads to extended lifespan; however, if raised at 15°C and shifted to 25°C at the L4 stage, worms are not long-lived (McElwee et al., 2004), all as in a similar *glp-1* mutant (Arantes-Oliveira et al., 2002). The latter, L4 shift treatment blocks germline development from L4 onwards, and is therefore expected to prevent tumor growth, and this was confirmed, and tumor suppression found to be unaffected by Carb (Figure 2D, Figure S2A). Notably, preventing bacterial proliferation caused *glp-4* to extend mean lifespan (Table S2): on Carb by +7.6% (*p* = 0.0012, Figure 2E), and on Kan by +11.8% (*p* = 0.022, Figure 2F). That a different intervention that blocks tumor development also extends lifespan when bacterial proliferation is prevented is further evidence for the unmasking hypothesis. However, it remains possible that *glp-4(bn2)* here extends lifespan by other mechanisms. Considering other senescent pathologies, *glp-4* (Carb present) in the main worsened them, on both D5 and D10 for gonadal degeneration and intestinal atrophy, though pharyngeal pathology was reduced on D10 (Figure S2B).

A further possibility is that *glp-4(bn2)* leads to a slight inhibition of signals emanating from the germline, whose life-extending effects are masked by bacterial infection. The life-extending effects of inhibiting germline development are dependent on the *daf-16* FOXO class forkhead transcription factor and the *daf-12* dafachronic acid receptor (Hsin and Kenyon, 1999; Motola et al., 2006). We therefore tested the capacity of *glp-4* to extend lifespan in *daf-16(mgDf50)* and *daf-12(m20)* mutant backgrounds. Both *daf-16* and *daf-12* alone moderately reduced lifespan at 25°C (-31.5% and -23.6%, respectively, *p* < 0.0001, *p* < 0.0001; Table S3, S4), consistent with prior observations (Larsen et al., 1995; Lin et al., 2001). Moreover, in the presence of Carb, *glp-4* again extended mean lifespan, by +8.4% (*daf-16* trials, *p* < 0.0001) or +9.1% (*daf-12* trials, *p* < 0.0001). Notably, in a *daf-16* background, *glp-4* increased mean lifespan by +21.5% (*p* < 0.0001), while in a *daf-12* background, it increased mean lifespan by +10.7% (*p* < 0.0001) (Figure 3A, B; Table S3, S4). Life extension by *glp-4(bn2)* (L4 shift, Carb present) is therefore not attributable to reduced germline signaling, consistent with a life-shortening effect of uterine tumor development.

**Figure 3.**
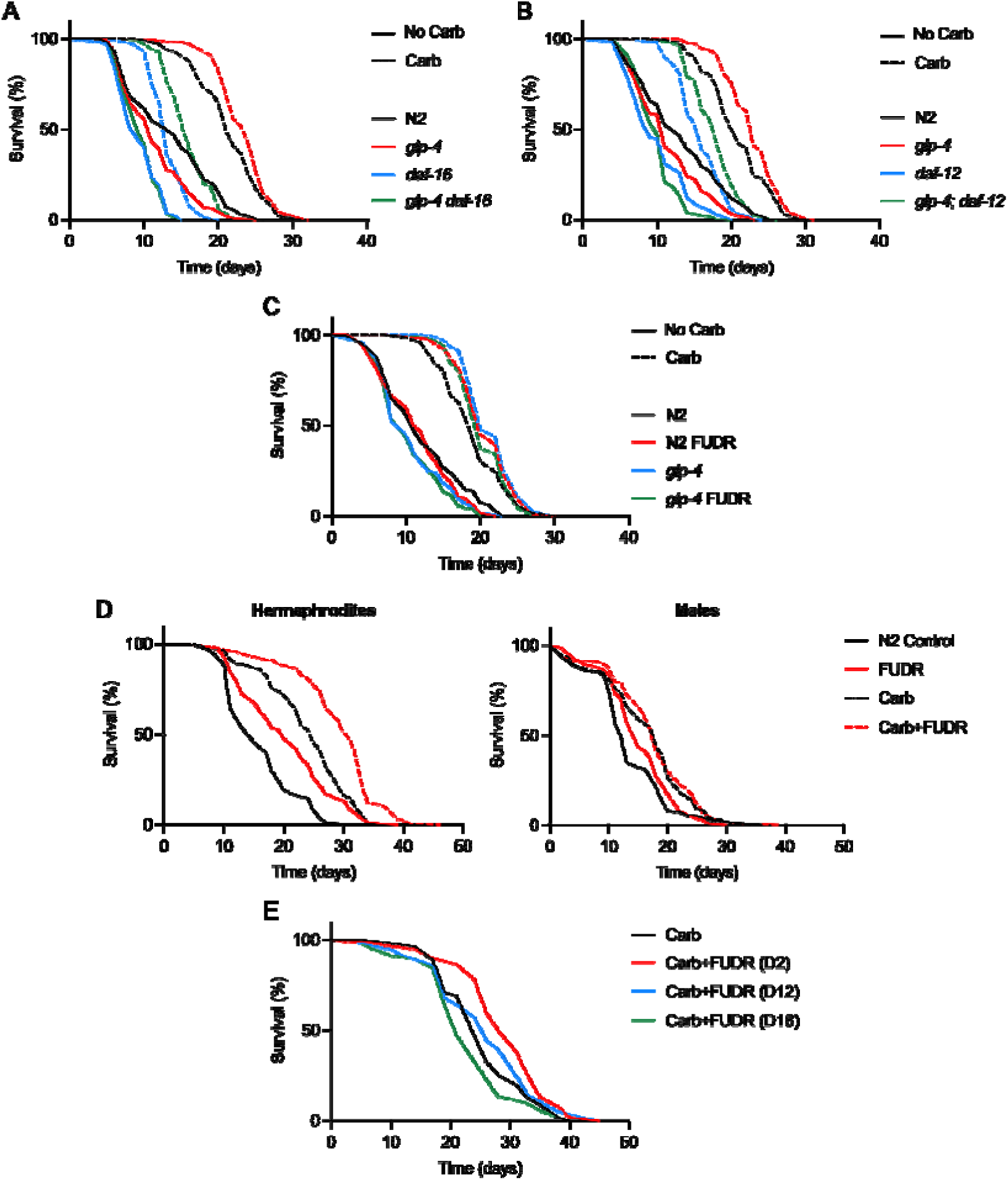
Evidence that uterine tumors shorten hermaphrodite lifespan when bacterial proliferation is suppressed. (A, B) Life extension by *glp-4(bn2)* is not (A) *daf-16* or (B) *daf-12* dependent (25°C). (C) FUDR does not increase lifespan in tumor-less *glp-4* mutant hermaphrodites. (D) On Carb, 50 μM FUDR increases lifespan of N2 hermaphrodites but not males (20°C). (E) FUDR treatment fails to extend lifespan when initiated after tumor development. For individual trials and statistical comparisons, see Tables S3 (A), S4 (B), S5 (C), S7 (D), and S8 (E).

### Further tests of the life-limiting tumor hypothesis

However, it remains possible that life-extending effects of FUDR and *glp-4* are due to changes other than tumor suppression. To test this further we used a triangulation strategy: the use of multiple, orthogonal tests to address a given question (Munafò and Davey Smith, 2018). First, we reasoned that if FUDR extends lifespan by suppressing tumor development then, in the presence of Carb, it should not extend lifespan in *glp-4* populations, since they are tumor-less. To test this, effects of 50 μM FUDR on N2 and *glp-4* populations were compared (L4 shift to 25°C, Carb present). FUDR increased lifespan in N2, again, but not in *glp-4* (+9.50%, *p* = 0.0032, -2.58%, *p* = 0.12, respectively; summed data; Figure 3C, Table S5).

We also checked that FUDR efficiently suppresses tumor development in N2 at 25°C. To our surprise, 50 μM and even 100 μM FUDR only marginally reduced tumor size (Figure S3A). This could partially reflect the fact that uterine tumors are smaller at 25°C (Figure S3B). Noting that FUDR only increased N2 lifespan (Carb present) in 2/3 trials (Table S5), we conducted 3 further trials to verify this. A significant lifespan increase was seen in only 1/3 cases (Table S6); combined data from all 6 trials indicates a significant 6.62% increase in mean lifespan after FUDR treatment (*p* = 0.0065; Table S6). We postulate that the smaller life-extending effect of FUDR at 25°C could reflect its weaker effect on tumor growth, and perhaps their being smaller in the absence of treatment (Figure S3B).

As a further test, we compared the effect of 50 μM FUDR on lifespan in hermaphrodites and males. Males, lacking either a uterus or the oocytes from which uterine tumors develop, do not develop uterine tumors (Kern et al., 2025). Thus if life extension by FUDR is attributable to tumor suppression, then this treatment should not extend male lifespan. To prevent life-shortening male-male interactions (Gems and Riddle, 2000) animals were cultured individually in liquid culture (McCulloch and Gems, 2003). In the presence of Carb addition of FUDR significantly increased lifespan in hermaphrodites (+22.0%, *p* < 0.0001) but not in males (+5.1%, *p* = 0.588) (Figure 3D, Table S7). Unexpectedly FUDR also increased lifespan in the absence of Carb, in both hermaphrodites (+27.5%, *p* < 0.0001) and males (+12.3%, *p* < 0.0027), suggesting the presence of different life-shortening mechanisms in liquid culture.

Next we tested the effect on lifespan of initiating exposure to FUDR at later ages, comparing initiation at D2, D12 and D18 of adulthood (Carb present). Our expectation was that treatment starting at the two later ages, after tumors have already formed, would not extend lifespan. This proved to be the case: FUDR from D2 increased lifespan, but from D12 and D18 did not (+14.28%, +1.65%, -11.80%, respectively, *p* = 0.0053, *p* = 0.36, *p* = 0.12, respectively, *N* = 1; Figure 3E, Table S8).

Finally we tested effects of FUDR on lifespan in other nematode species, comparing two sibling species pairs where one species is androdioecious (hermaphrodites [H] and males) and the other gonochoristic (females [F] and males). Aging *C. tropicalis* and *Pristionchus pacificus* (hermaphrodites) develop uterine tumors while females of their respective sibling species, *C. wallacei* and *P. exspectatus*, do not (Kern et al., 2023). If FUDR extends lifespan by preventing tumor growth, then it should extend lifespan in species with hermaphrodites but not females.

On Carb, FUDR increased lifespan in *C. tropicalis* hermaphrodites but also in *C. wallacei* females (+20.7%, +13.3%, respectively, *p* = 0.0022, *p* = 0.046; Figure S4A, Table S9), and the effect in the former was not significantly greater than in the latter (*p* = 0.181, Cox Proportional Hazard [CPH] analysis). On Carb, FUDR increased lifespan in *P. pacificus* hermaphrodites but decreased it in *P. exspectatus* females (+7.1%, -15.6%, respectively, *p* = 0.0001, *p* < 0.0001; Figure S4B, Table S10). Thus, findings from only the *Pristionchus* sibling species pair support the view that FUDR extends lifespan by inhibiting tumor development.

Taken together, with the exception of the *Caenorhabditis* sibling species pair tests, results of these additional tests support the view that the life-extending effects of FUDR and *glp-4* in the presence of Carb are attributable to tumor suppression.

### Uterine tumors partially mask the life-shortening effect of vitellogenesis

Next we explored the position of other causes and types of pathology in the hierarchy of death. The *C. elegans* intestine is the site of synthesis of yolk, which is transported across the body cavity to provision developing oocytes (Kimble, 1983; Perez and Lehner, 2019). Vitellogenins (yolk proteins) are the products of the genes *vit-1* - *vit-6*, and RNAi knockdown of *vit* expression blocks vitellogenin accumulation, reduces intestinal atrophy and yolk pool accumulation, and extends lifespan (Ezcurra et al., 2018; Murphy et al., 2003; Sornda et al., 2019). To test the possibility that the effect on late-life mortality of blocking yolk synthesis is partially masked by either bacterial infection or uterine tumor development, we subjected N2 populations to *vit-5,-6* RNAi, in the presence of either the antibiotic kanamycin (Kan), or FUDR, or both. Kan was used because the plasmid in the *E. coli* RNAi feeding strains confers Carb resistance. To exclude the possibility that Kan interferes with RNAi knockdown (given its impact on the *E. coli*), effects of *gfp* RNAi on a transgenic *Pftn-1::GFP* strain (Ackerman and Gems, 2012) was compared in the absence and presence of Kan, but no effect of Kan on RNAi knockdown of GFP was detected (Figure S5A).

We first examined effects on pathology. Tumor size was reduced by FUDR under all conditions (± Kan, ± RNAi) on D7 but, unexpectedly, not on D14 (Figure 4A; for all statistical comparisons, see Table S11), in contrast to prior tests (Figure 2B). This may reflect the use here of the *E. coli* RNAi feeding strain HT115, rather than OP50 as previously. This was confirmed using a direct test (Figure 4B): suppression of tumor growth by FUDR was greater on *E. coli* OP50 than HT115, possibly reflecting differences in FUDR biotransformation (Scott et al., 2017).

**Figure 4.**
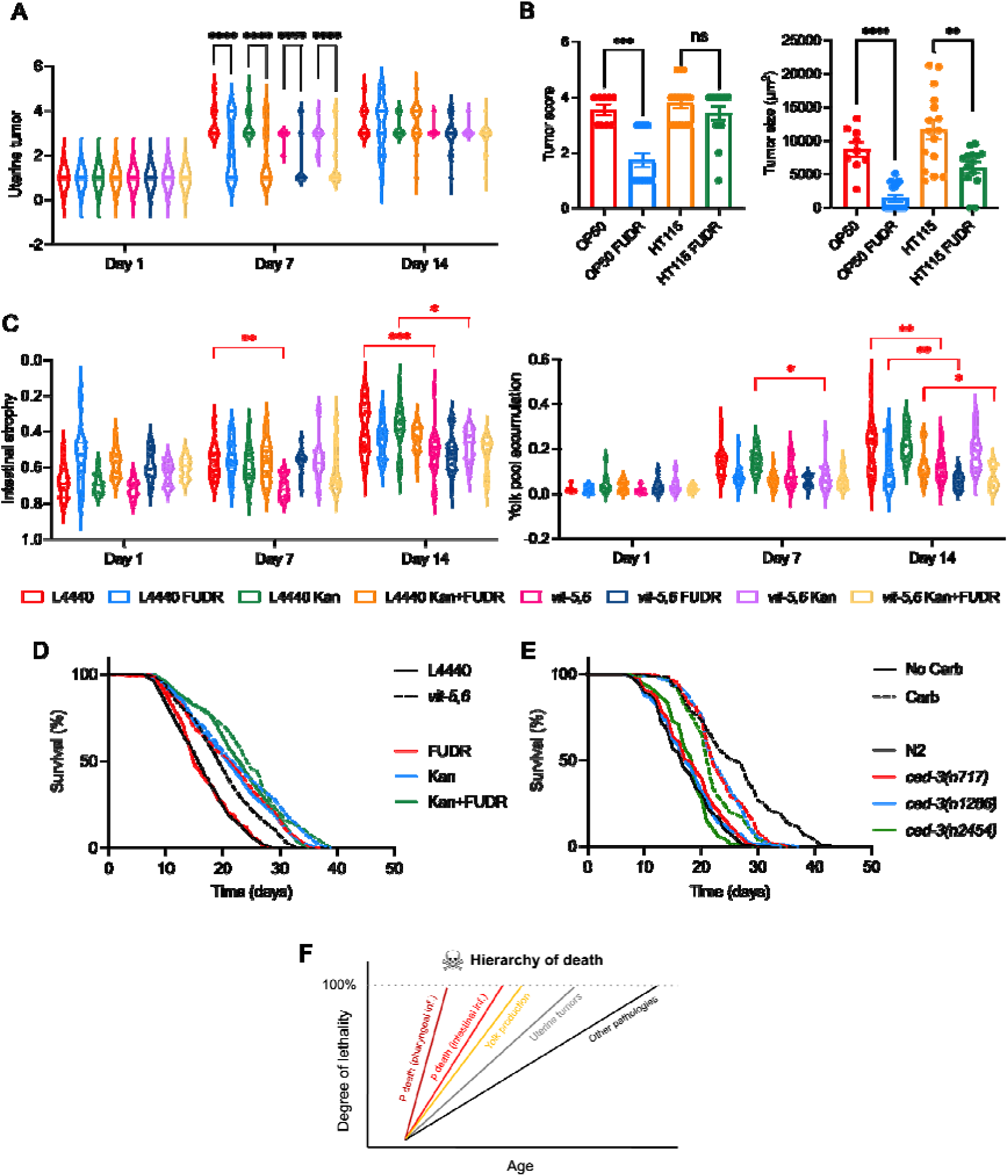
Condition-dependent effects of *vit-5,-6* RNAi and *ced-3(n717)*. (A) Suppression of tumor growth by FUDR on D7 but not D14 under all conditions. (B) Weaker tumor suppression by 50 μM FUDR on *E. coli* HT115 than OP50 (D8 of adulthood). Error bars, standard error. (C) *vit-5,-6* RNAi reduced intestinal atrophy (left) and yolk pool accumulation (right). (D) Effects of *vit-5,-6* RNAi on lifespan ± FUDR ± Kan. (A-D) Trials performed with FUDR from L4 onwards. Effects on lifespan of FUDR from L4 showed some variability, such that in earlier trials, life extension was seen with Carb present. We were unable to identify the cause of this variability. (E) Effects of *ced-3* on lifespan ± Carb. For individual trials and statistical comparisons in lifespan analyses, see Tables S11 (D) and S13 (E). (F) Hierarchy of death competing risks (onion) model. The evidence presented clearly supports the view that death from infection masks life-limiting effects of uterine tumors. The weaker life-extending effects of blocking yolk synthesis when Kan is present suggests an approximate position for vitellogenesis higher up the hierarchy than uterine tumors.

*vit-5,-6* RNAi by itself reduced intestinal atrophy and yolk pool accumulation (Figure 4C, Table S11), as previously seen (Ezcurra et al., 2018; Sornda et al., 2019). Kan alone did not significantly reduce pathologies, and on Kan or FUDR rescuing effects of *vit-5,-6* RNAi were still detectable (Figure 4C, Table S11). However, at some ages FUDR alone reduced yolk pool size (Figure 4C), and also pharyngeal and distal gonad pathology (Figure S5B).

Next, effects on lifespan were examined, first considering those of *vit-5,-6* RNAi. In the absence of Kan or FUDR, *vit-5,-6* RNAi extended mean lifespan (+20.4%, *p* < 0.0001, Figure 4B; Table S12), as previously seen (Sornda et al., 2019). Notably, life extension by *vit-5,-6* RNAi was smaller on Kan (+5.34%, *p* = 0.019; Figure 4B; Table S12), though comparing effects of *vit-5,-6* RNAi with and without Kan, the difference in this diminution did not quite reach statistical significance (*p* = 0.0659, CPH analysis). Thus, in contrast to uterine tumor development, prevention of bacterial infection does not unmask a greater life-shortening effect of vitellogenesis. The apparent reduction in the life-extending effect caused by Kan could imply that *vit-5,-6* RNAi increases infection resistance, e.g. by suppressing intestinal atrophy.

The impact of FUDR on life extension by *vit-5,-6* RNAi was also assessed, first in the absence of Kan. Notably, the presence of FUDR increased the life-extending effect of *vit-5,-6* RNAi to +28.2% (*p* < 0.0001 log rank test; CPH, *p* = 0.0133; Figure 4B; Table S12). Thus, blocking vitellogenesis causes uterine tumors to become life-limiting, even in the presence of proliferative bacteria. This again suggests that *vit-5,-6* RNAi increases infection resistance. Consistent with this, in the presence of Kan and FUDR, the life-extending effect of *vit-5,-6* RNAi is again reduced, to +5.38% (*p* = 0.014 log rank test; CPH, *p* = 0.0241; Figure 4B; Table S12).Taken together, these results suggest that the life-extending effect of *vit-5,-6* RNAi is attributable to increased resistance to late-life *E. coli* infection. In the presence of such protective changes, uterine tumors contribute to late-life mortality, such that their suppression enhances the life-extending effect of *vit-5,-6* RNAi.

### Delayed distal gonad degeneration does not extend lifespan in the absence of infection

During aging of *C. elegans* hermaphrodites, the distal gonad undergoes marked atrophy and then fragments (Figure 1B) (de la Guardia et al., 2016; Garigan et al., 2002). Notably, this pathology was reduced by FUDR, significantly so in the absence of Carb (Figure S1B; D14 of adulthood). One contributor to such atrophy is futile, post-reproductive continuation of physiological apoptosis in the distal gonad, which in young adults contributes to oogenesis (Gumienny et al., 1999). Supporting this, prevention of physiological apoptosis using *ced-3* mutations, including *ced-3(n717)*, which block both somatic and germline apoptosis, significantly delays gonad degeneration (de la Guardia et al., 2016). Prior reports of effects of inhibition of *ced-3* on lifespan are conflicting, for reasons unknown, describing either no effects using *ced-3(n1286*) (Garigan et al., 2002), *ced-3(n717)* or *ced-3* RNAi (de la Guardia et al., 2016), or increased lifespan using *ced-3(n717*) (Judy et al., 2013) or *ced-3* RNAi (Curran and Ruvkun, 2007).

To test the possibility that effects of distal gonad degeneration on late-life mortality are masked by death due to infection, lifespan of N2 and *ced-3* populations were compared in the absence or presence of Carb. Three *ced-3* mutants were tested, *ced-3(n717)*, *ced-3(n1286)* and *ced-3(n2454)*, but no extension of lifespan relative to N2 controls was seen under either condition (Figure 4E; Table S13). This suggests that distal gonad disintegration does not become life-limiting in the absence of infection. In fact, preventing bacterial infection unmasked a life-shortening effect of *ced-3* (-10.8% to -16.1%, *p* < 0.0001; Figure 4C; Table S13).

### *glp-4* extends *daf-2* lifespan by increasing infection resistance

*daf-2* insulin/IGF-1 signaling (IIS) mutants are long-lived (Kenyon et al., 1993) and exhibit increased resistance to bacterial infection (Garsin et al., 2003; Podshivalova et al., 2017; Troemel et al., 2006; Zhao et al., 2021). *daf-2* also strongly suppresses several senescent pathologies, but only weakly reduces uterine tumor size (Ezcurra et al., 2018), though it does substantially reduce tumor DNA content (Golden et al., 2007; McGee et al., 2012). We noted a previous observation that *glp-4(bn2)* (L4 temperature upshift) extends lifespan in *daf-2* mutant backgrounds (no Carb), including the class 1 (less pleiotropic) *daf-2(m577)* mutant (Gems et al., 1998; McElwee et al., 2004). This could imply that, as with *vit-5,-6* RNAi treatment (see above), *daf-2* mutant infection resistance causes uterine tumors to become life limiting.

To investigate this we re-examined the effect of *glp-4* on lifespan in a *daf-2(m577)* background (no Carb). This confirmed that *glp-4* extends *daf-2* mean lifespan (+61.0%, *p* < 0.0001; Figure S6A; Table S14), as previously reported (McElwee et al., 2004). If *glp-4* enhances *daf-2* longevity by preventing tumor formation, then this enhancing effect should still be present in the presence of Carb. However, this proved not to be the case: *glp-4* only marginally increased lifespan in a *daf-2(m577)* background (+0.45%, *p* = 0.0015, Figure S6B, Table S14).

*glp-4* enhancement of infection resistance could involve prevention of either of the two forms of infection related death in *C. elegans*. These are the P (“big P”) deaths with a swollen, infected pharynx, and the later p (“small p”) deaths with an atrophied pharynx (Zhao et al., 2017). Necropsy analysis comparing *daf-2* and *glp-4; daf-2* populations showed a modest reduction in P death frequency, and strong increases in both P and p lifespan (Figure S6C-E, Table S15; *N* = 1). This implies that in a *daf-2* background *glp-4* enhances resistance to both pharyngeal and intestinal infection.

These results suggest the presence of a weak reduction in germline signaling in *glp-4* (L4 shift) animals, that in a *daf-2*(+) background is not sufficient to increase lifespan, but which synergises with the reduction in IIS in *daf-2* mutants, leading to increased infection resistance and, consequently, increased lifespan. They also suggest that uterine tumors do not contribute to late-life mortality in *daf-2* mutants (at least in class 1 mutants).

## Discussion

This study deals with the question of how to interpret the effects of treatments that extend lifespan in model organism studies. A standard interpretation, explicit or implicit, is that an increase in lifespan represents a slowing of the overall aging process, unless it is clearly the result of some form of environmental disruption, as in the case of life-shortening *E. coli* infection in *C. elegans*. Here we show that a senescent pathology of intrinsic and quasi-programmed origin, uterine tumors, can contribute to late-life mortality. Notably, this contribution is condition dependent: under standard laboratory conditions (with proliferating *E. coli*) it is masked, presumably by lethal late-life infection.

This represents a distant mirror of human mortality: in centuries past, the major causes of death were infectious pathogens (e.g. tuberculosis, smallpox, plague), made more lethal by malnutrition, cold and poor hygiene. Thanks to improvements in conditions in the developed world, the main causes of death are now cardiovascular disease, cancer and lung disease (particularly chronic obstructive pulmonary disease). Hypothetically, a cure for all cancer would cause a greater increase in lifespan in the 21st than in the 14th century; similarly, preventing uterine tumor development extends *C. elegans* lifespan in the absence but not presence of bacterial infection.

### Mapping out the hierarchy of death

These findings draw attention to the utility of viewing *C. elegans* lifespan as the product of competing causes of mortality. Competing risks approaches have been used in studies of human mortality rate (Austin et al., 2016). Such competing causes of mortality are a possible determinative factor in the outcome of tests of interactions between interventions that extend *C. elegans* lifespan, that can be difficult to interpret (Gems et al., 2002). For example, *rsks-1(ok1255)*, affecting S6 kinase in the mTOR pathway, modestly increases mean lifespan (+20%), but in a *daf-2(e1370)* background it causes a massive ∼360% increase relative to *daf-2* alone (Chen et al., 2013; Selman et al., 2009). This could imply, in a competing risks interpretation, that IIS more than mTOR signaling limits wild-type lifespan, but reducing IIS unmasks a greater life-limiting role for mTOR.

Thus, interpretation of *C. elegans* lifespan data would be aided by a description of the competing causes of late-life mortality, both extrinsic and intrinsic. Part of such a description is an account of how such competing causes mask one another, and their position in an onion-like, nested hierarchy of causes. Here we present the beginnings of such an account (Figure 4F).

The outer-most layer of the onion, in the presence of proliferating *E. coli*, is death with pharyngeal infection (P death). Here death from pharyngeal infection is the outer skin of the onion for P but not p individuals. This was demonstrated by an earlier study, in which mutation of *eat-2*, which reduces pharyngeal pumping rate, led to reduction of P death frequency (Zhao et al., 2017). The resulting increase in p deaths, which occur later, caused an increase in overall lifespan. The lifespan of the p subpopulation, like that of the P subpopulation, is limited by *E. coli* infection, since preventing the latter increases p lifespan (Zhao et al., 2017). Given the findings in the present study, we can confidently state that uterine tumor development becomes life-limiting when bacterial infection is prevented. Key evidence for this is the life-extending effects of two interventions that prevent tumor formation, FUDR and *glp-4(bn2)*, but only when bacterial proliferation is prevented (Figure 2), and only in comparison to control strains in which tumors are present (Figure 2, 3).

Our analysis allowed additional pathologies to be placed, tentatively, in the hierarchy of death (Figure 4F). Blocking vitellogenesis by *vit* gene RNAi modestly increases lifespan (Murphy et al., 2003; Sornda et al., 2019), i.e. the presence of proliferating *E. coli* does not mask an effect on lifespan; however, it remained possible that partial masking occurs. In fact, antibiotic treatment reduced rather than increased the life-extending effect of inhibiting vitellogenesis (Figure 4D). This implies that life-extension here is largely due to increased infection resistance, placing vitellogenesis above tumor development in the hierarchy of death (Figure 4F).

Distal gonad atrophy and fragmentation is an anatomically striking pathology, yet its inhibition by mutation of *ced-3* did not extend lifespan, even in the presence of Carb (Figure 4E). This could imply either that distal gonad atrophy never contributes to mortality, or that its effects on mortality are masked by competing risks. A further consideration is that *ced-3* delays but does not prevent distal gonad atrophy, to which declining germline stem cell proliferation likely also contributes (de la Guardia et al., 2016; Luo et al., 2010); thus, full suppression of distal gonad atrophy might reveal a contribution to late-life mortality.

### Uterine tumors as a distant mirror of cancer

*C. elegans* uterine tumor is a form of age-related neoplasia that in certain respects resemble its mammalian counterparts. First, it has etiological and phenotypic similarities to mammalian teratomas, particularly ovarian cysts (Wang et al., 2018b). Second, like mammalian neoplasia, it appears largely as part of organismal senescence, and can contribute to late-life mortality (this study). Third, *C. elegans* uterine tumor can be treated with FUDR, a drug also used for the treatment of colorectal cancer in humans (Allen-Mersh et al., 1994). These similarities support the presence of underlying, general principles of senescent pathophysiology, that are operative across the animal kingdom during aging, from nematodes to humans (Ezcurra et al., 2018; Gems and de Magalhães, 2021). Hence we may learn about such general principles from studying organisms even as primitive as *C. elegans*. It is worth noting too that a screen for compounds that prevent uterine tumor growth could have identified FUDR, a useful anticancer drug. Thus, the capacity to prevent *C. elegans* uterine tumor growth could in principle be used to identify novel anti-cancer agents. A further deduction is that if a given drug extends lifespan in *C. elegans* in the absence but not the presence of proliferating *E. coli*, this could reflect action through suppression of uterine tumor growth.

Also of interest is the mechanism by which uterine tumors contribute to late-life mortality, which could be informative with respect to how cancer causes illness and death. Uterine tumors often press against and flatten the intestine, providing one potential mechanism. A second possibility is that uterine tumors secrete molecular species (e.g. proteins) that have deleterious effects on other tissues; the *C. elegans* uterus does contain secreted proteins whose levels increase with age and shorten life (Zimmerman et al., 2015). A third is that uterine tumors absorb vital nutrients in a sponge-like fashion, leading to health-impairing deficiencies in other tissues; consistent with this, uterine tumors accumulate vitellogenin (Wang et al., 2018b). Any such effects are likely to entail an interaction with age-related frailty due to other aspects of organismal aging.

More broadly, this study serves as a reminder that interventions that extend lifespan in model organisms may not be assumed to affect the aging process as a whole.

## Materials and methods

### Culture methods and strains

*C. elegans* maintenance was performed using standard protocols (Brenner, 1974). Except where noted, all strains were grown at 20°C on agar plates containing nematode growth media (NGM, containing Bacto Peptone [Gibco, USA]), seeded with *E. coli* OP50 as a food source. In certain experiments (as specified), bacterial lawns were treated an antibiotic (4 mM carbenicillin, unless otherwise stated) to inhibit bacterial infection of *C. elegans*.

An N2 hermaphrodite stock recently obtained from the Caenorhabditis Genetics Center was used as wild type (N2H) (Zhao et al., 2019). Other *C. elegans* strains used included: DR20 *daf-12(m20) X*, DR1567 *daf-2(m577) III*, GA134 *glp-4(bn2) I; daf-2(m577) III*, GA633 *wuIs177*[*Pftn-1::GFP, lin-15(+)*] *daf-2(m577) III*, GA1952 *daf-16(mgDf50) I*, GA6001 *glp-4(bn2) I*; *daf-12(m20) X*, GA6002 *glp-4(bn2) daf-16(mgDf50) I,* MT1522 *ced-3(n717) IV*, MT3002 *ced-3(n1286) IV,* MT8354 *ced-3(n2454) IV,* SS104 *glp-4(bn2) I*. Other nematode species strains (wild type): JU1373 *C. tropicalis*, JU1873 *C. wallacei*, PS312 *Pristionchus pacificus*, RS5522 *Pristionchus exspectatus*.

### Survival analysis

Nematodes were maintained at a density of 25-30 per plate, and transferred daily during the egg laying period, followed by every 6-7 days thereafter. The L4 stage was defined as day 0. Mortality was scored every 1-2 days, with worms scored as alive if they showed any movement, spontaneously or in response to gentle touch with a worm pick. Raw data for all lifespan trials in this study is presented in Dataset S1.

Due to high frequency of plate leaving by solitary males, it is difficult to maintain them on agar plates. To circumvent this, lifespan measurements in trials including males were performed in liquid medium. Animals were cultured singly in 96-well microtitre plates, in 50 μL of a suspension of *E. coli* in S medium (concentration 1 × 10^8^ −1 × 10^9^ cells mL^−1^), as described (McCulloch and Gems, 2003).

### Mortality deconvolution analysis

This analysis is based on the presence of two forms of death in aging *C. elegans* cultured on proliferating *E. coli*: earlier death with an infected, swollen pharynx (P death) and later death with an atrophied pharynx (p death) (Zhao et al., 2017). Mortality deconvolution involves analysis of P and p lifespans separately. Alterations in lifespan can result from altered percentages of P (and p) deaths, and/or altered P and/or p lifespan. Deconvolved mortality statistics include each of these values. Corpses were scored by necropsy as P or p using the highest magnification of a dissecting microscope, as described (Zhao et al., 2017).

### Treatment with FUDR

NGM plates were seeded with *E. coli* OP50, and left overnight at room temperature for lawns to grow. FUDR was then added using a 5 mM stock to give a final concentration of 50 μM. FUDR treatment was initiated on D2 of adulthood unless otherwise stated.

### RNA-mediated interference (RNAi)

*E. coli* HT115 RNAi-producing and control bacteria (plasmid L4440), were cultivated as previously described (Kamath et al., 2003), seeded onto NGM plates containing carbenicillin and IPTG. IPTG was added to cultured bacteria and induced at 37°C for 1 hr before seeding plates. RNAi treatment was started at the L4 stage, after larval development on HT115 (no carbenicillin).

### Analysis of senescent pathology

This was performed as previously described (Ezcurra et al., 2018; Kern et al., 2025). At each age assayed, 10-15 animals were mounted onto 2% agar pads and anesthetized with 0.2% levamisole. Images were acquired using differential interference contrast (DIC, or Nomarski) optics, with a Zeiss ApoTome.2 microscope. A Hamamatsu C13440 ORCA-Flash4.0 V3 digital camera and Zen software were used for image acquisition. For pharynx, distal gonad and tumor pathologies, images were examined by trained scorers, assigned severity scores of 1-5, and mean values calculated. Scoring criteria were: 1 = youthful, healthy appearance; 2 = early signs of mild deterioration; 3 = clearly discernible, mild pathology; 4 = well-developed pathology; and 5 = very severe pathology (e.g. gonad completely disintegrated), or reaching a maximal level (e.g. large uterine tumor filling the entire body cavity in the mid-body region). Intestinal atrophy was quantified by measuring the intestinal width in a region posterior to the uterine tumors, subtracting the width of the intestinal lumen and dividing by the body width. Yolk accumulation was measured by dividing the yolk pool area by the area of the body visible in the field of view as captured at 630x magnification.

### Statistical analysis

Statistical tests were performed on raw data using GraphPad Prism 9.0 (GraphPad Software, USA) and JMP Pro 15 (JMP Statistical Discovery LLC, USA) unless otherwise stated, with the specific tests and post hoc corrections performed as described in the figure legends. No statistical methods were used to predetermine sample size. The experiments were not randomized. The investigators were not blinded to allocation during experiments and outcome assessment.

## Acknowledgments

We thank R.J. Sommer (M.P.I. Developmental Biology, Tübingen) for providing *Pristionchus* species, Lucy Guo, Siu Sylvia Lee (Cornell) and Bruce Zhang for useful discussion and/or comments on the manuscript, and Xinyu Cheng, Anna Girtle, James Rawson, Zihe Wang and Yihan Wu for minor research contributions. Some strains were provided by the Caenorhabditis Genetics Center, which is funded by NIH Office of Research Infrastructure Programs (P40 OD010440). D.G. is grateful for the support of a Wellcome Trust Investigator Award (215574/Z/19/Z).

## Author contributions

Conceptualization, D.G.; methodology, C.C.K., D.G., H.W.; investigation, A.S.A., M.E., Y.F., C.C.K., C.N.H., J.Q., H.W., A.Z.; writing – original draft, D.G.; writing – review and editing, D.G., H.W.; funding acquisition, D.G.; supervision, D.G.

## Supplementary Information

### Other supplementary files

**Supplementary Dataset 1.** All raw lifespan data.

## Supplementary figures

**Supplementary Figure 1.**
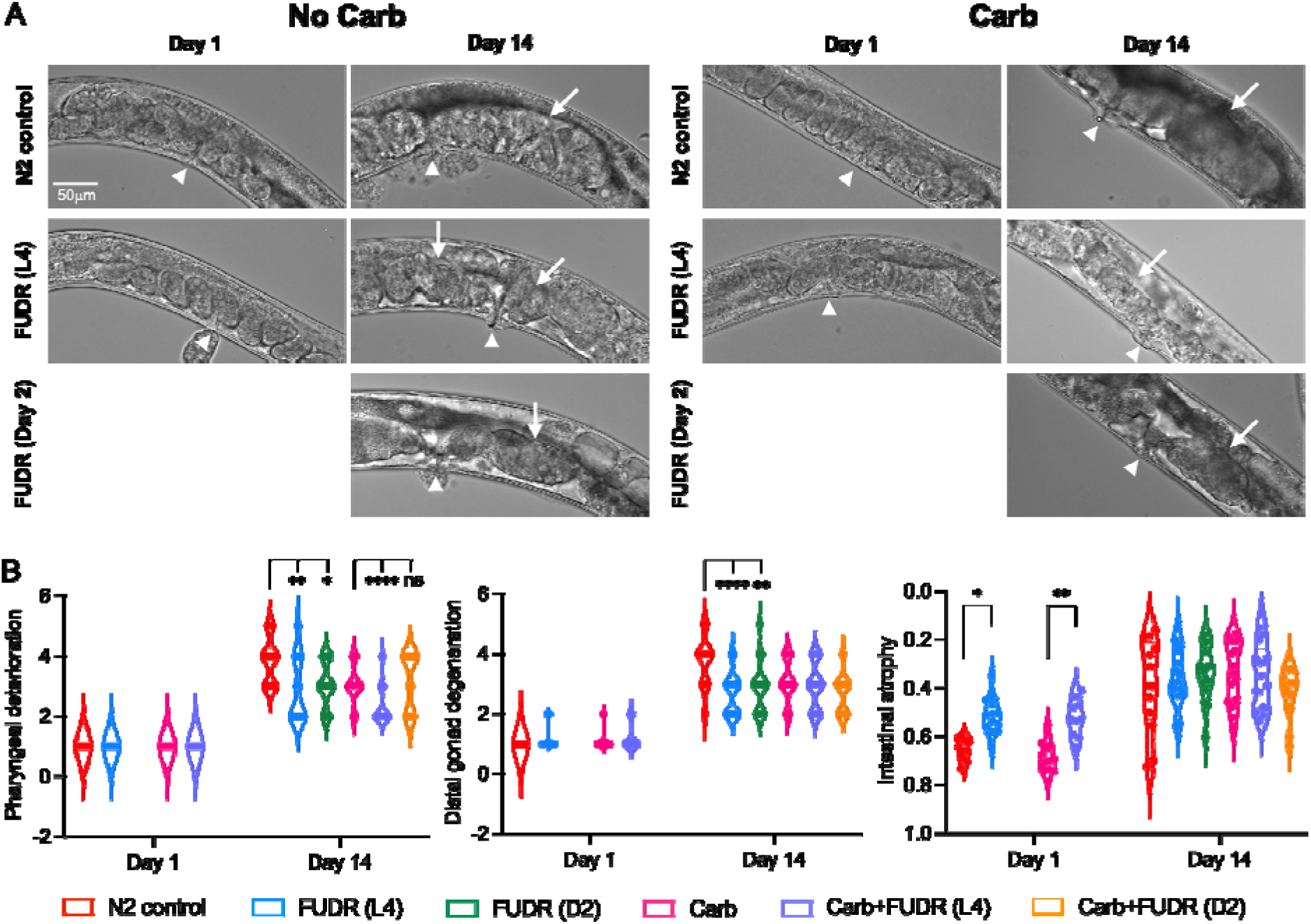
Effects of FUDR on senescent pathologies in *C. elegans* hermaphrodites (20°C). (**A**) Representative images of young and old nematodes. Arrows, uterine tumors; arrowheads, vulvae. (**B**) Effect of FUDR on three senescent pathologies. The apparent increase in intestinal atrophy present on D1 with FUDR from L4 likely reflects inhibition of intestinal growth.

**Supplementary Figure 2.**
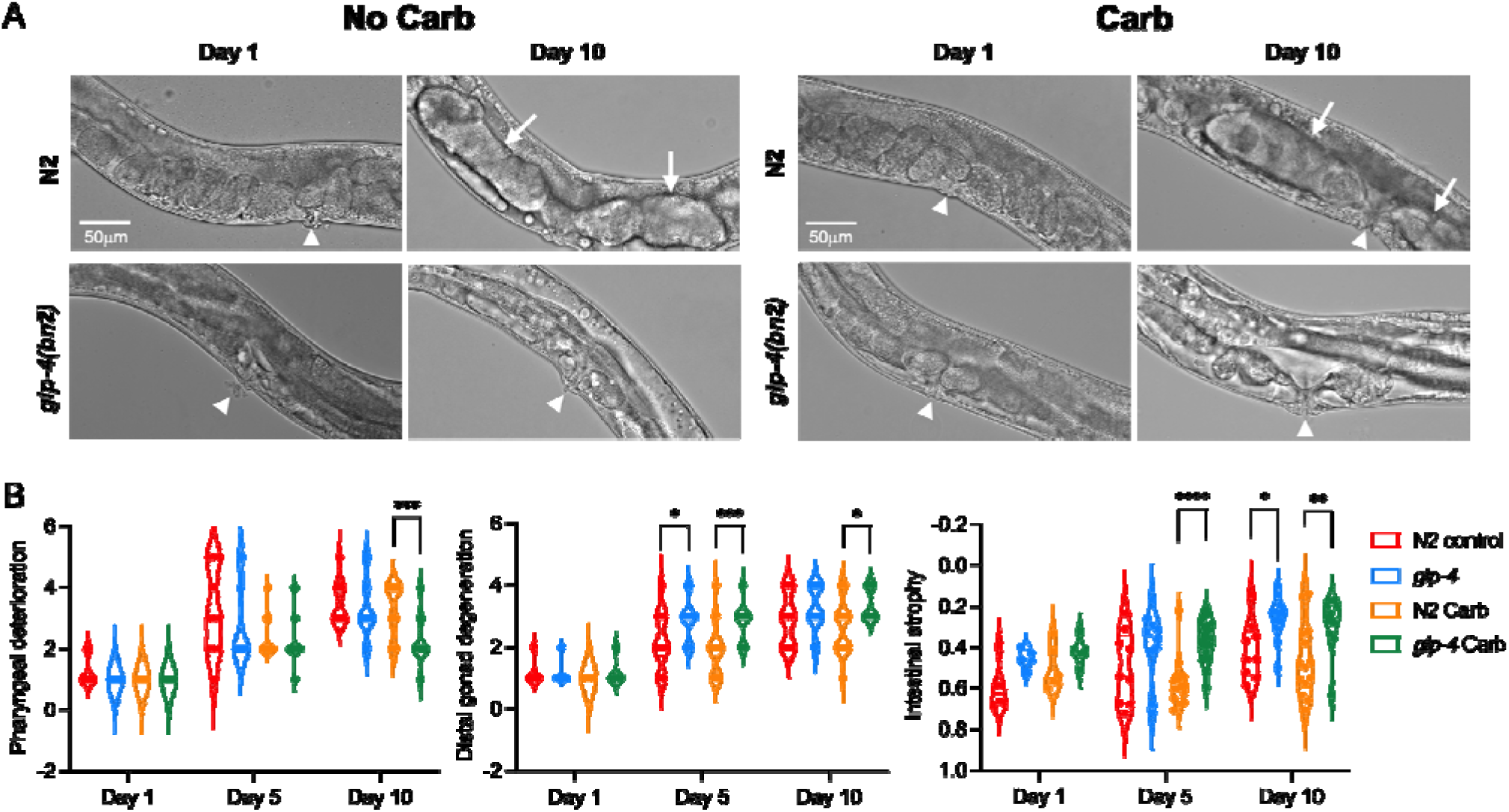
Effects of *glp-4(bn2)* on aging pathologies (25°C from L4). (**A**) Representative images of young and old nematodes. Arrows, uterine tumors (absent in old *glp-4* animal); arrowheads, vulvae. (**B**) Effects of *glp-4(bn2)* and Carb on three senescent pathologies.

**Supplementary Figure 3.**
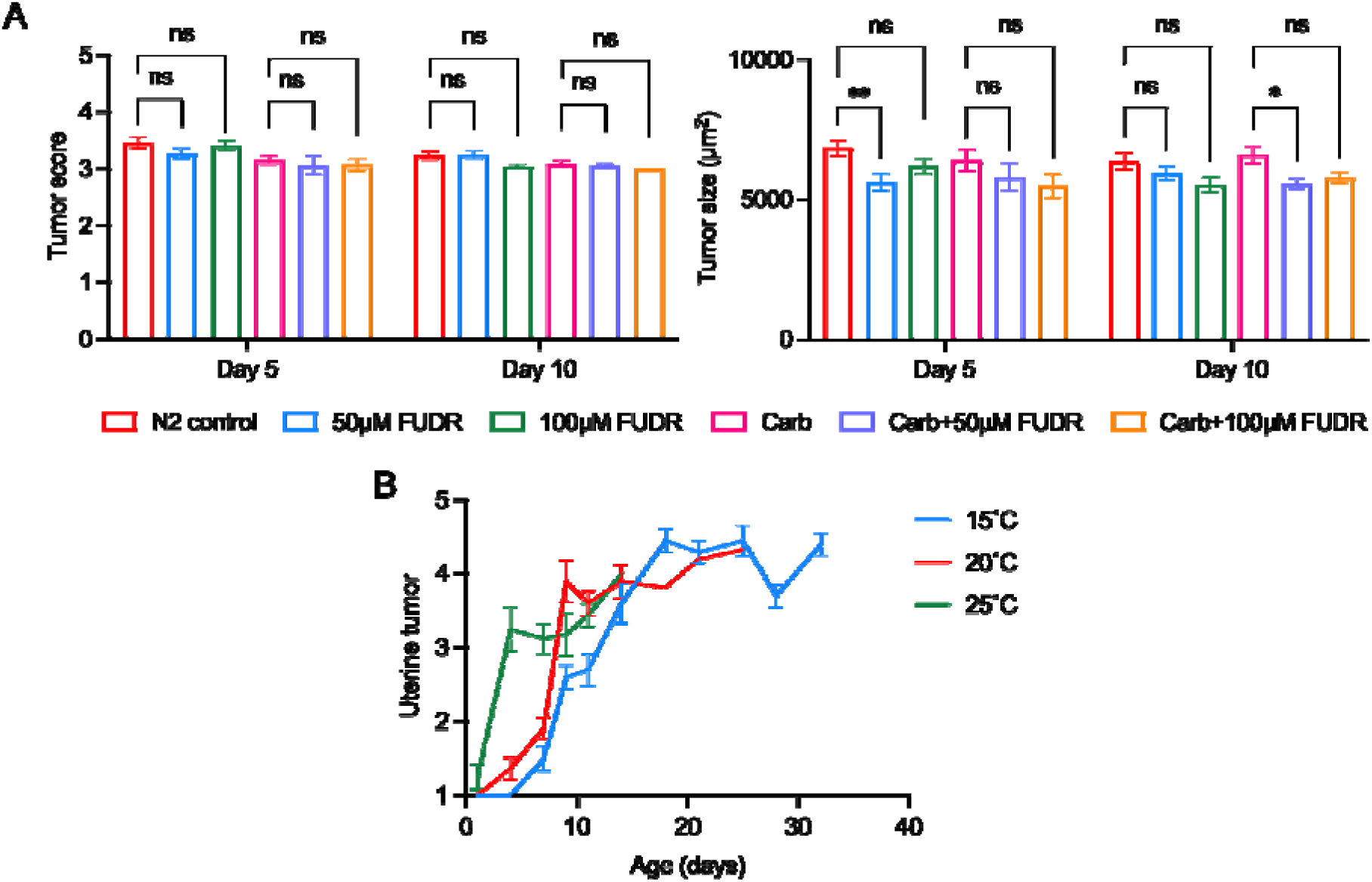
Effects of FUDR on lifespan and uterine tumor development (25°C). (**A**) FUDR does not increase lifespan in tumor-less *glp-4* mutant hermaphrodites. (**B**) Development of uterine tumor under different temperatures (15°C, 20°C, 25°C). (C) At 25°C, 50μM, 100μM FUDR only marginally reduce tumor size; tumor scale (left), tumor size (right). Measurement of tumor size (cross sectional area) is more accurate than the uterine tumor score.

**Supplementary Figure 4.**
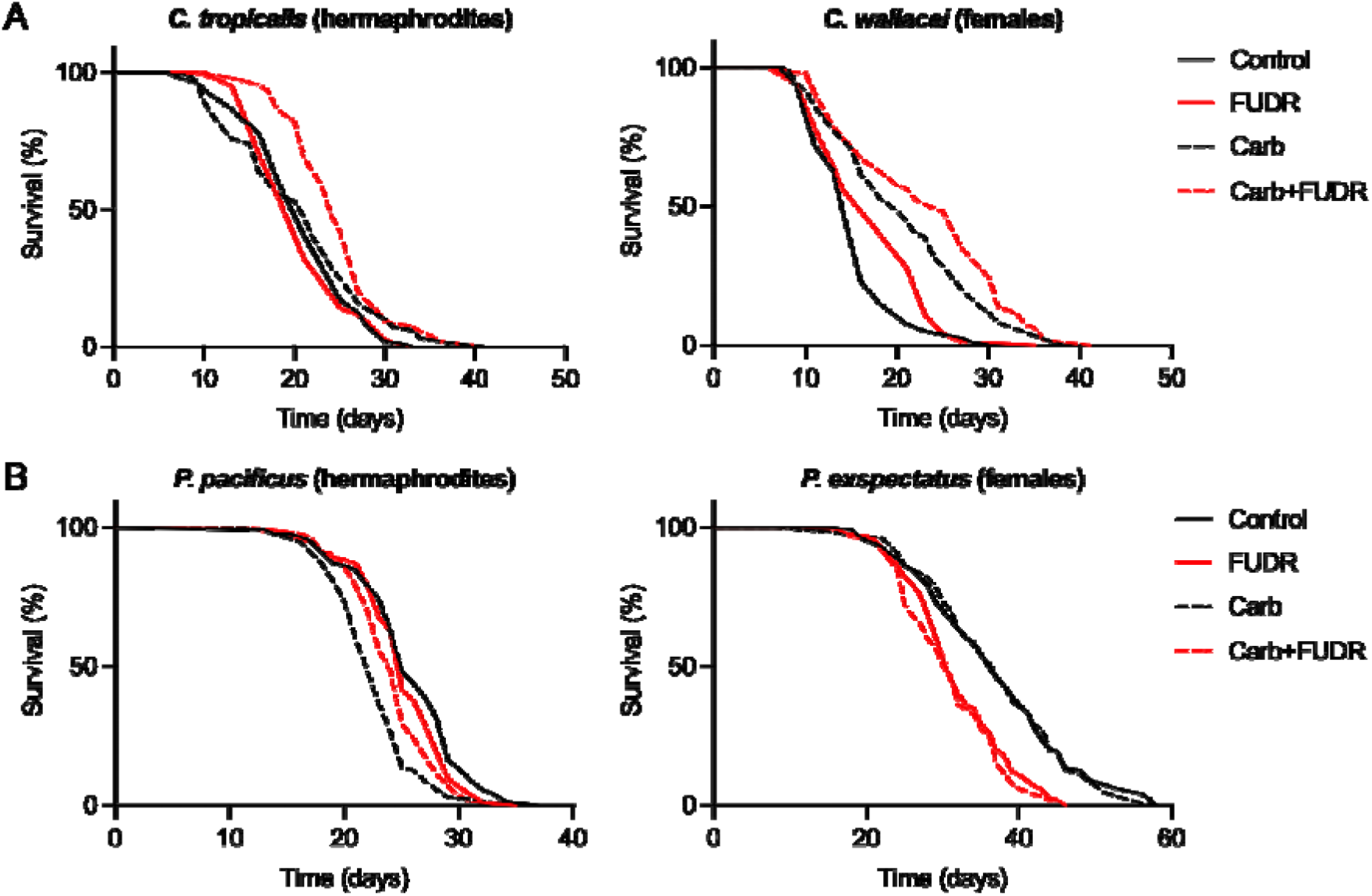
Effects of FUDR on hermaphrodites and females of two sibling species pairs. (**A**) *Caenorhabditis tropicalis* (hermaphrodites) and *C. wallacei* (females). (**B**) *Pristionchus pacificus* (hermaphrodites) and *P. exspectatus* (females).

**Supplementary Figure 5.**
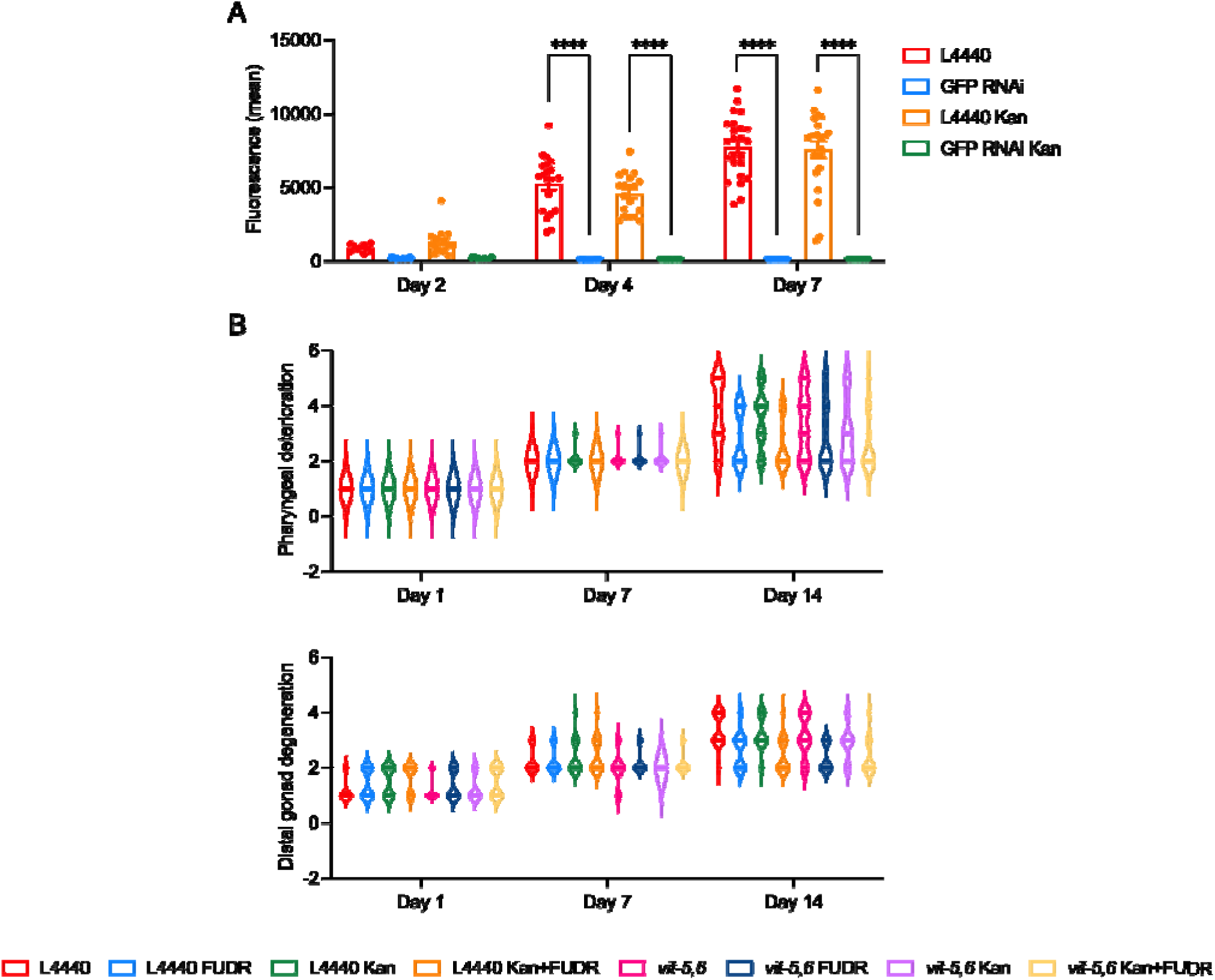
Effects of treatment combinations to remove life-limiting pathologies. (**A**) No effect of Kan on knockdown of *pftn-1::gfp* by *gfp* RNAi. Error bars: standard error. (**B**) Effects of treatment combinations on pharyngeal deterioration and distal gonad degeneration.

**Supplementary Figure 6.**
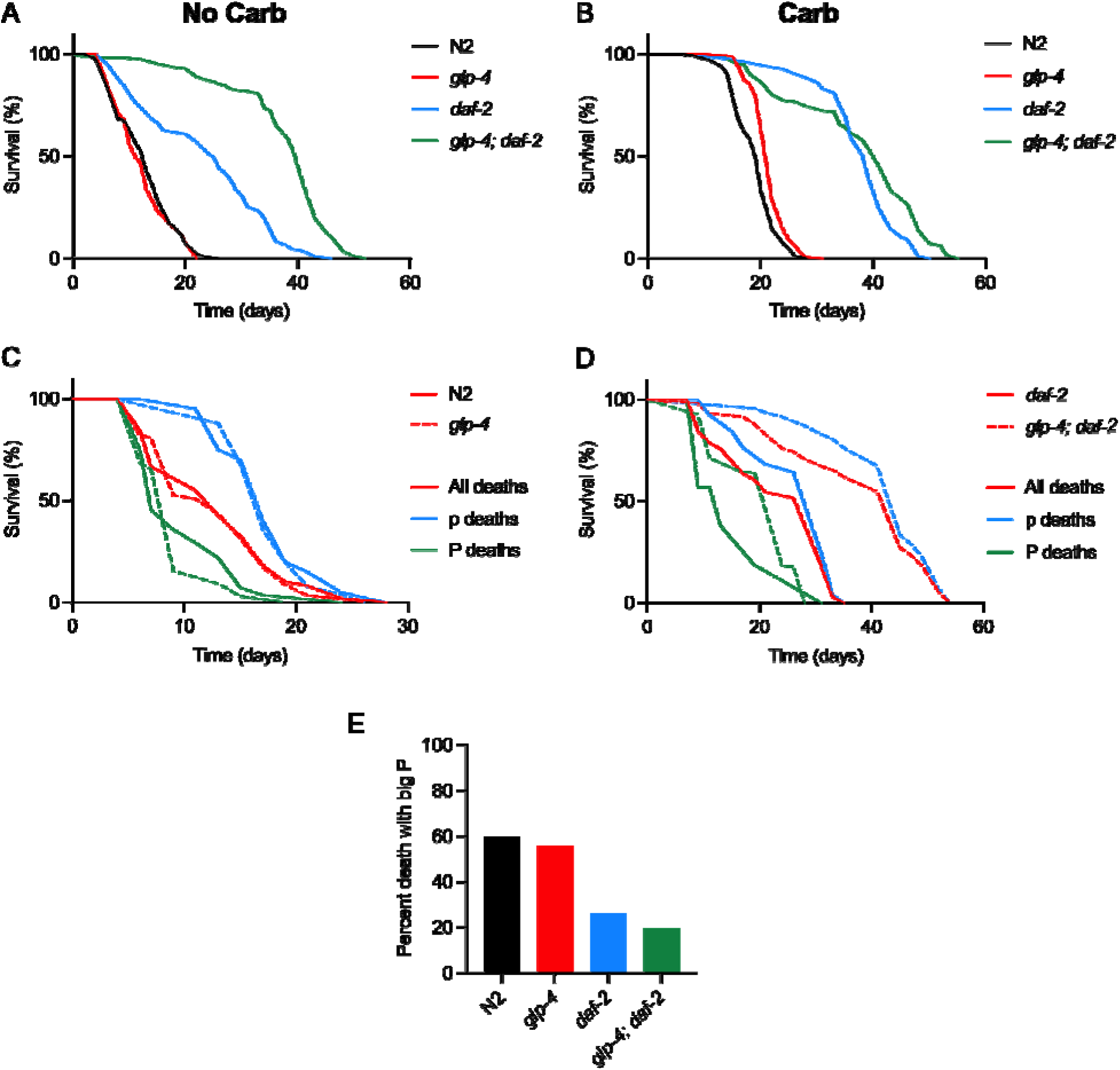
*glp-4(bn2)* increases *daf-2(m577)* lifespan by increasing infection resistance (25°C from L4 stage). (**A**, **B**) *glp-4* increases *daf-2(m577)* longevity in the absence (**A**) but not presence (**B**) of Carb. (**C**, **D**) Mortality deconvolution analysis of effects of *glp-4* and *daf-**2***. Effects of *glp-4* in a wild-type (**C**) and *daf-2(m577)* (**D**) background (no Carb). (**E**) P death frequency is strongly reduced by *daf-2* but not *glp-4*.

**Table S1.** Effects of 50 mM FUDR on lifespan in the presence of Carb or Kan (20°C)

| Strain/<br>conditions | Number<br>of deaths/<br>censored | Mean<br>[median]<br>lifespan<br>(days) | % change<br>vs.<br>N2 control | p vs. N2<br>(log rank) | % change<br>vs.<br>non FUDR | p vs. non<br>FUDR<br>(log rank) |
| --- | --- | --- | --- | --- | --- | --- |
| N2 control | [C] 153/27<br>[1] 54/8<br>[2] 51/9<br>[3] 50/10 | 16.82 [16]<br>17.31 [15]<br>17.29 [16]<br>15.82 [17] |  |  |  |  |
| N2<br>FUDR (from L4) | [C] 167/13<br>[1] 56/4<br>[2] 57/3<br>[3] 54/6 | 15.79 [15]<br>16.20 [15]<br>15.88 [16]<br>15.28 [14] | -6.12 [-6.25]<br>-6.41 [0]<br>-8.16 [0]<br>-3.41 [-17.65] | 0.0024<br>0.0945<br>0.0397<br>0.1805 | -6.12 [-6.25]<br>-6.41 [0]<br>-8.16 [0]<br>-3.41 [-17.65] | 0.0024<br>0.0945<br>0.0397<br>0.1805 |
| N2<br>FUDR (from D2) | [C] 161/19<br>[1] 51/9<br>[2] 55/5<br>[3] 55/5 | 17.84 [19]<br>18.88 [21]<br>17.65 [18]<br>17.07 [17] | +6.06 [+18.75]<br>+9.07 [+40.00]<br>+2.08 [+12.50]<br>+7.90 [0] | 0.2742<br>0.2787<br>0.9850<br>0.3516 | +6.06 [+18.75]<br>+9.07 [+40.00]<br>+2.08 [+12.50]<br>+7.90 [0] | 0.2742<br>0.2787<br>0.9850<br>0.3516 |
| N2 Carb | [C] 148/22<br>[1] 40/10<br>[2] 52/8<br>[3] 56/4 | 25.61 [25]<br>23.50 [21]<br>24.54 [25]<br>28.11 [28] | +52.26 [+56.25]<br>+35.76 [+40.00]<br>+41.93 [+56.25]<br>+77.69 [+64.71] | <0.0001<br><0.0001<br><0.0001<br><0.0001 |  |  |
| N2<br>Carb + FUDR<br>(from L4) | [C] 166/14<br>[1] 53/7<br>[2] 55/5<br>[3] 58/2 | 24.12 [24]<br>24.70 [25]<br>23.20 [23]<br>24.47 [24] | +43.40 [+50.00]<br>+42.69 [+66.67]<br>+34.18 [+43.75]<br>+54.68 [+41.18] | <0.0001<br><0.0001<br><0.0001<br><0.0001 | -5.82 [-4.00]<br>+5.11 [+19.05]<br>-5.46 [-8.00]<br>-12.95 [-14.29] | <0.0001<br>0.9100<br>0.0267<br><0.0001 |
| N2<br>Carb + FUDR<br>(from D2) | [C] 163/17<br>[1] 52/8<br>[2] 56/4<br>[3] 55/5 | 31.25 [32]<br>26.92 [27]<br>33.75 [35]<br>32.80 [33] | +85.79 [+100.00]<br>+55.52 [+80.00]<br>+95.20 [+118.75]<br>+107.33 [+94.12] | <0.0001<br><0.0001<br><0.0001<br><0.0001 | +22.02 [+28.00]<br>+14.55 [+28.57]<br>+37.53 [+40.00]<br>+16.68 [+17.86] | <0.0001<br>0.0359<br><0.0001<br>0.0035 |
| N2 Kan | [C] 156/24<br>[1] 46/14<br>[2] 56/4<br>[3] 54/6 | 25.13 [25]<br>22.87 [23]<br>27.32 [25]<br>24.80 [24] | +49.41 [+56.25]<br>+32.12 [+53.33]<br>+58.01 [+56.25]<br>+56.76 [+41.18] | <0.0001<br><0.0001<br><0.0001<br><0.0001 |  |  |
| N2 Kan + FUDR<br>(from L4) | [C] 161/19<br>[1] 51/9<br>[2] 52/8<br>[3] 58/2 | 23.75 [24]<br>24.76 [25]<br>23.83 [25]<br>22.78 [24] | +41.20 [+50.00]<br>+43.04 [+66.67]<br>+37.83 [+56.25]<br>+43.99 [+41.18] | <0.0001<br><0.0001<br><0.0001<br><0.0001 | -5.49 [-4.00]<br>+8.26 [+8.70]<br>-12.77 [0]<br>-8.15 [0] | <0.0001<br>0.7003<br>0.0011<br>0.0085 |
| N2 Kan + FUDR<br>(from D2) | [C] 141/39<br>[1] 44/16<br>[2] 52/8<br>[3] 45/15 | 29.61 [30]<br>28.84 [30]<br>29.63 [30]<br>30.33 [31] | +76.04 [+87.50]<br>+66.61 [+100.00]<br>+71.37 [+87.50]<br>+91.72 [+82.35] | <0.0001<br><0.0001<br><0.0001<br><0.0001 | +17.83 [+20.00]<br>+26.10 [+30.43]<br>+8.46 [+20.00]<br>+22.30 [+29.17] | <0.0001<br><0.0001<br>0.0779<br>0.0019 |

**Table S2.**
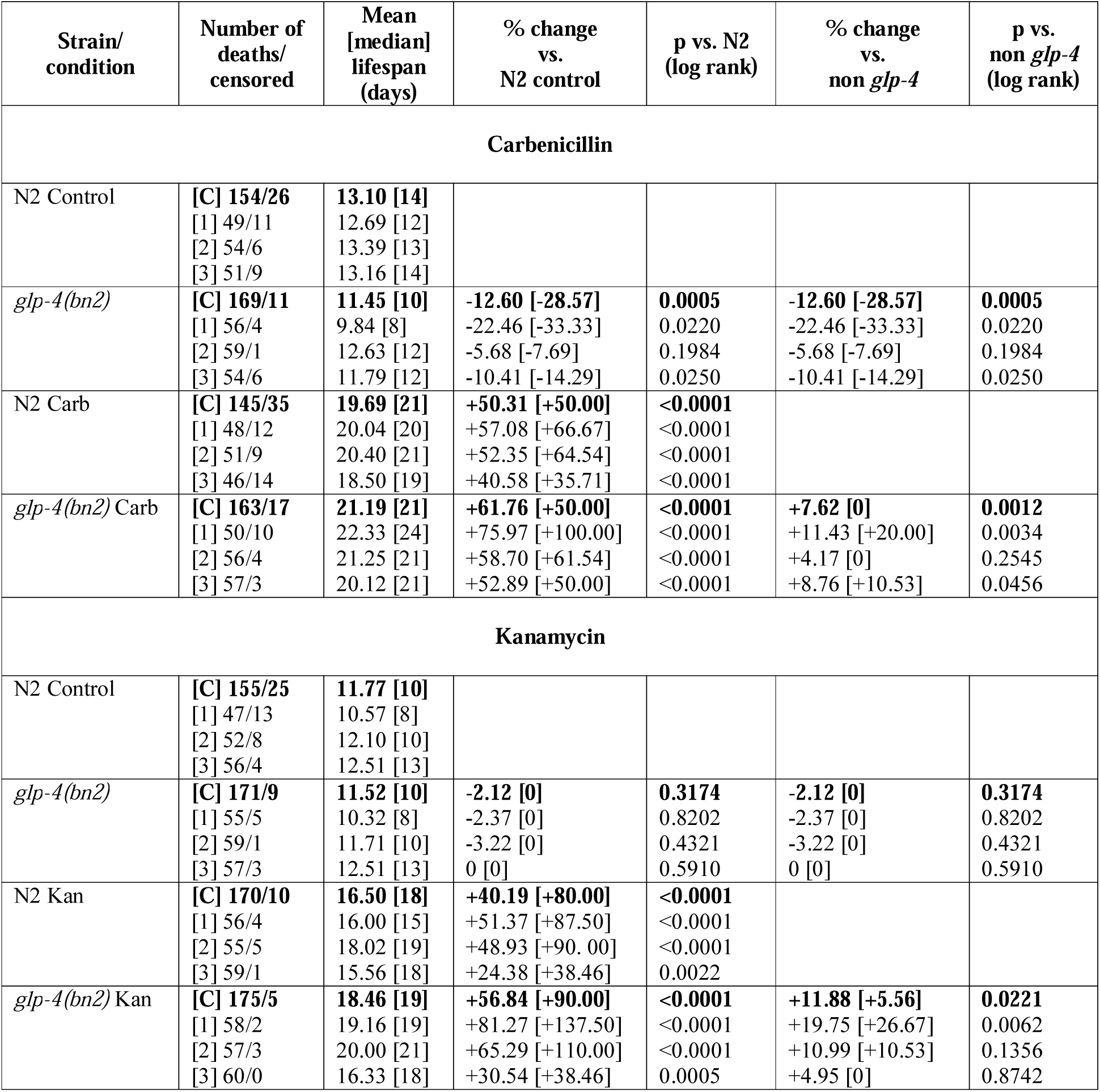
Effects of *glp-4(bn2)* on lifespan in the presence of Carb or Kan (25°C from L4 stage)

**Table S3.**
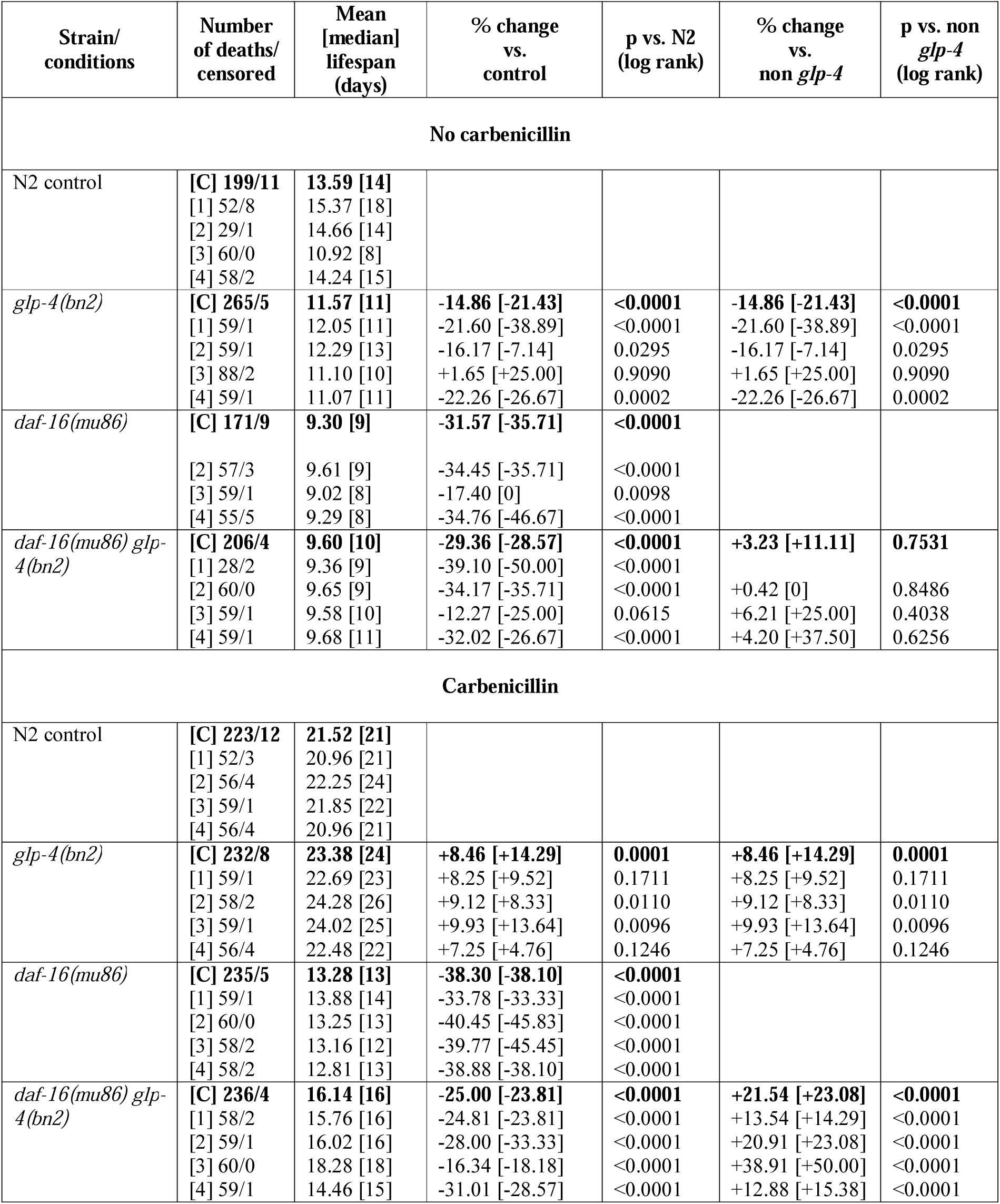
Effects of *daf-16* on *glp-4* longevity (25°C from L4)

**Table S4.**
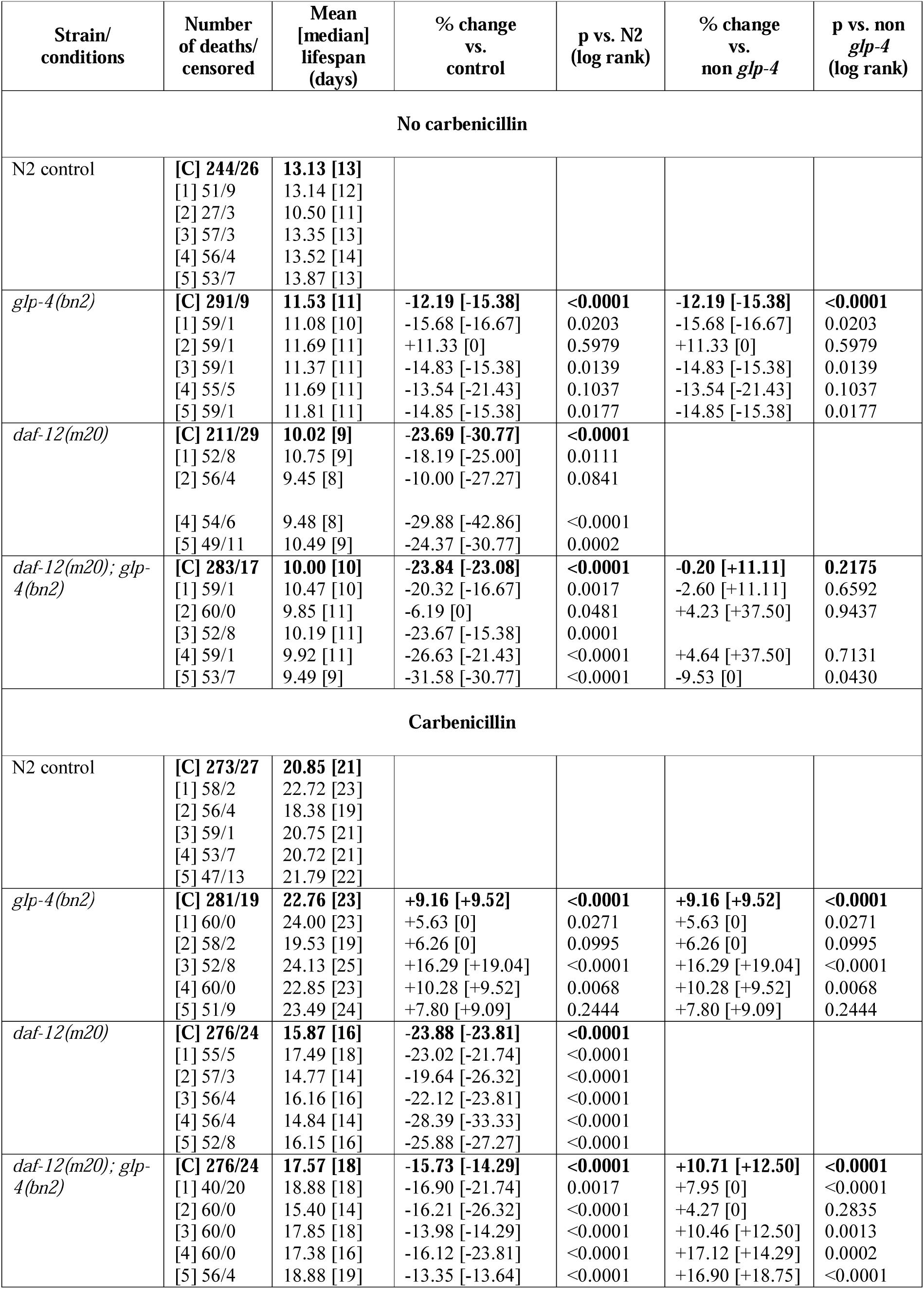
Effects of *daf-12* on *glp-4* longevity (25°C from L4)

**Table S5.** Effects of FUDR on *glp-4(bn2)* lifespan (25°C from L4 stage)

| Strain/<br>condition | Number of<br>deaths/<br>censored | Mean<br>[median]<br>lifespan (days) | % change<br>vs.<br>N2 control | p vs.<br>N2 control<br>(log rank) | % change<br>vs.<br>*non glp-4* | p vs.<br>*non glp-4*<br>(log rank) | % change<br>vs.<br>untreated | p vs.<br>untreated (log<br>rank) |
| --- | --- | --- | --- | --- | --- | --- | --- | --- |
| **No carbenicillin** |  |  |  |  |  |  |  |  |
| N2 control | [C] **114/16**<br>[1] 26/4<br>[2] 40/0<br>[3] 48/12 | **12.14 [11]**<br>11.87 [11]<br>11.95 [11]<br>12.38 [12] |  |  |  |  |  |  |
| N2 FUDR (D2) | [C] **139/21**<br>[1] 52/8<br>[2] 40/0<br>[3] 47/13 | **11.70 [12]**<br>11.74 [13]<br>11.68 [12]<br>11.71 [12] | **-3.62 [+9.09]**<br>-1.10 [+18.18]<br>-2.26 [9.09]<br>-5.41 [0] | **0.1672**<br>0.6440<br>0.4558<br>0.4450 |  |  | **-3.62 [+9.09]**<br>-1.10 [+18.18]<br>-2.26 [9.09]<br>-5.41 [0] | **0.1672**<br>0.6440<br>0.4558<br>0.4450 |
| *glp-4(bn2)* | [C] **177/3**<br>[1] 60/0<br>[2] 60/0<br>[3] 57/3 | **10.78 [10]**<br>10.62 [8]<br>10.55 [8]<br>11.19 [10] | **-11.20 [-9.09]**<br>-10.53 [-27.27]<br>-11.72 [-27.27]<br>-9.61 [-16.67] | **0.0129**<br>0.2372<br>0.1610<br>0.1061 | **-11.20 [-9.09]**<br>-10.53 [-27.27]<br>-11.72 [-27.27]<br>-9.61 [-16.67] | **0.0129**<br>0.2372<br>0.1610<br>0.1061 |  |  |
| *glp-4(bn2)*<br>FUDR (D2) | [C] **179/1**<br>[1] 59/1<br>[2] 60/0<br>[3] 60/0 | **10.48 [10]**<br>9.86 [8]<br>11.05 [11]<br>10.52 [10] | **-13.67 [-9.09]**<br>-16.93 [-27.27]<br>-7.53 [0]<br>-15.02 [-16.67] | **0.0006**<br>0.0473<br>0.1839<br>0.0143 | **-10.43 [-16.67]**<br>-16.01 [-38.46]<br>-5.39 [-8.33]<br>-10.16 [-16.67] | **0.0169**<br>0.0218<br>0.5738<br>0.0847 | **-2.78 [0]**<br>-7.16 [0]<br>+4.74 [+37.50]<br>-5.99 [0] | **0.3262**<br>0.2510<br>0.8902<br>0.3383 |
| **Carbenicillin** |  |  |  |  |  |  |  |  |
| N2 Control | [C] **143/17**<br>[1] 39/1<br>[2] 45/15<br>[3] 59/1 | **19.15 [20]**<br>18.97 [18]<br>19.71 [20]<br>18.85 [20] |  |  |  |  |  |  |
| N2<br>FUDR (D2) | [C] **132/18**<br>[1] 37/3<br>[2] 44/6<br>[3] 51/9 | **20.97 [20]**<br>21.17 [23]<br>20.43 [20]<br>21.27 [20] | **+9.50 [0]**<br>+11.60 [+27.78]<br>+3.65 [0]<br>+12.84 [0] | **0.0032**<br>0.0399<br>0.7487<br>0.0104 |  |  | **+9.50 [0]**<br>+11.60 [+27.78]<br>+3.65 [0]<br>+12.84 [0] | **0.0032**<br>0.0399<br>0.7487<br>0.0104 |
| *glp-4(bn2)* | [C] **170/10**<br>[1] 58/2<br>[2] 59/1<br>[3] 53/7 | **21.57 [20]**<br>21.03 [20]<br>22.03 [23]<br>21.66 [20] | **+12.64 [0]**<br>+10.86 [+11.11]<br>+11.77 [+15.00]<br>+14.91 [0] | **<0.0001**<br>0.0500<br>0.0240<br>0.0021 | **+12.64 [0]**<br>+10.86 [+11.11]<br>+11.77 [+15.00]<br>+14.91 [0] | **<0.0001**<br>0.0500<br>0.0240<br>0.0021 |  |  |
| *glp-4(bn2)*<br>FUDR (D2) | [C] **178/2**<br>[1] 60/0<br>[2] 60/0 | 20.43 [20]<br>20.05 [20]<br>21.27 [23] | **+6.68 [0]**<br>+5.69 [+11.11]<br>+7.91 [+15.00] | **0.0615**<br>0.4077<br>0.2421 | **-2.58 [0]**<br>-5.29 [-13.04]<br>+4.11 [+15.00] | **0.1203**<br>0.0601<br>0.3464 | **-5.29 [0]**<br>-4.66 [0]<br>-3.45 [0] | **0.0031**<br>0.1408<br>0.1595 |
|  | [3] 58/2 | 19.97 [20] | +5.94 [0] | 0.2855 | -6.11 [0] | 0.0929 | -7.80 [0] | 0.0242 |

**Table S6.** Verification of effects of FUDR on *glp-4(bn2)* lifespan (25°C from L4 stage)

| Strain/<br>conditions | Number<br>of deaths/<br>censored | Mean<br>[median]<br>lifespan (days) | % change<br>vs.<br>N2 Carb | p vs. N2 Carb<br>(log rank) |
| --- | --- | --- | --- | --- |
| N2 Carb | [C] 303/22<br>[1] 39/1<br>[2] 45/15<br>[3] 59/1<br>[4] 59/1<br>[5] 58/2<br>[6] 43/2 | **18.14 [18]**<br>18.97 [18]<br>19.71 [20]<br>18.85 [20]<br>20.58 [21]<br>15.36 [16]<br>15.19 [14] |  |  |
| N2<br>Carb + FUDR (D2) | [C] 312/18<br>[1] 37/3<br>[2] 44/6<br>[3] 51/9<br>[4] 60/0<br>[5] 60/0<br>[6] 60/0 | **19.34 [19]**<br>21.17 [23]<br>20.43 [20]<br>21.27 [20]<br>21.08 [21]<br>16.38 [16]<br>17.02 [16] | **+6.62 [+5.56]**<br>+11.60 [+27.78]<br>+3.65 [0]<br>+12.84 [0]<br>+2.43 [0]<br>+6.64 [0]<br>+12.05 [+14.29] | **0.0065**<br>0.0399<br>0.7487<br>0.0104<br>0.3715<br>0.1037<br>0.0103 |

**Table S7.** Effects of 50 mM FUDR on lifespan of N2 hermaphrodites and males in monoxenic liquid culture (20°C, monoxenic liquid culture).

| Strain/<br>condition | Number of<br>deaths/<br>censored | Mean<br>[median]<br>lifespan<br>(days) | % change<br>vs.<br>N2 control | *p* vs.<br>N2 control<br>(log rank) | % change<br>vs.<br>untreated | *p* vs.<br>untreated<br>(log rank) |
| --- | --- | --- | --- | --- | --- | --- |
| *C. elegans* (hermaphrodites) |  |  |  |  |  |  |
| N2 control | [C] 13/131<br>[1] 5/43<br>[2] 6/42<br>[3] 2/46 | 15.95<sup>1</sup> [15]<br>13.98 [11]<br>16.30 [15]<br>17.59 [15] |  |  |  |  |
| N2 FUDR (D2) | [C] 3/141<br>[1] 3/45<br>[2] 0/48<br>[3] 0/48 | 20.34 [20]<br>18.56 [18]<br>20.77 [22]<br>21.56 [22] | +27.52 [+33.33]<br>+32.76 [+63.64]<br>+27.42 [+46.67]<br>+22.57 [+46.67] | <0.0001<br><0.0001<br>0.0010<br>0.0042 | +27.52 [+33.33]<br>+32.76 [+63.64]<br>+27.42 [+46.67]<br>+22.57 [+46.67] | <0.0001<br><0.0001<br>0.0010<br>0.0042 |
| N2 Carb | [C] 22/122<br>[1] 9/39<br>[2] 8/40<br>[3] 5/43 | 23.91 [25]<br>22.62 [23]<br>23.91 [26]<br>25.07 [25] | +49.91 [+66.67]<br>+61.80 [+109.09]<br>+46.69 [+73.33]<br>+42.52 [+66.67] | <0.0001<br><0.0001<br><0.0001<br><0.0001 |  |  |
| N2 Carb FUDR<br>(D2) | [C] 19/125<br>[1] 7/41<br>[2] 1/47<br>[3] 11/37 | 29.19 [31]<br>27.93 [30]<br>29.82 [33]<br>29.91 [32] | +83.01[+106.67]<br>+99.79 [+172.72]<br>+82.94 [+120.00]<br>+70.04 [+113.33] | <0.0001<br><0.0001<br><0.0001<br><0.0001 | +22.08 [+24.00]<br>+23.47 [+30.43]<br>+24.72 [+26.92]<br>+19.31 [+28.00] | <0.0001<br><0.0001<br><0.0001<br>0.0012 |
| *C. elegans* (males) |  |  |  |  |  |  |
| N2 control | [C] 5/139<br>[1] 4/44<br>[2] 0/48<br>[3] 1/47 | 13.48<sup>1</sup> [12]<br>12.53 [13]<br>12.71 [12]<br>15.24 [11] |  |  |  |  |
| N2 FUDR (D2) | [C] 4/140<br>[1] 1/47<br>[2] 2/46<br>[3] 1/47 | 15.14 [15]<br>12.04 [11]<br>16.84 [17]<br>16.56 [15] | +12.31 [+25.00]<br>-3.91 [-15.38]<br>+32.49 [+41.67]<br>+8.66 [+36.36] | 0.0027<br>0.5591<br><0.0001<br>0.8250 | +12.31 [+25.00]<br>-3.91 [-15.38]<br>+32.49 [+41.67]<br>+8.66 [+36.36] | 0.0027<br>0.5591<br><0.0001<br>0.8250 |
| N2 Carb | [C] 16/128<br>[1] 5/43<br>[2] 11/37<br>[3] 0/48 | 16.60 [18]<br>15.11 [16]<br>15.35 [17]<br>19.29 [20] | +23.15 [+50.00]<br>+20.59 [+23.08]<br>+20.77 [+41.67]<br>+26.57 [+81.82] | 0.0001<br>0.0591<br>0.0049<br>0.1031 |  |  |
| N2 Carb FUDR<br>(D2) | [C] 9/136<br>[1] 6/42<br>[2] 1/47<br>[3] 2/46 | 17.45 [18]<br>14.50 [16]<br>17.13 [17]<br>20.53 [22] | +29.45 [+50.00]<br>+15.72 [+23.08]<br>+34.78 [+41.67]<br>+34.71 [+100.00] | <0.0001<br>0.2128<br>0.0001<br>0.0088 | +5.12 [0]<br>-4.04 [0]<br>+11.60 [0]<br>+6.43 [+10] | 0.5883<br>0.2413<br>0.1236<br>0.1362 |

<sup>1</sup>Note that in none of the three trials do solitary males live longer than solitary hermaphrodites. This is surprising given earlier studies that observed that solitary N2 males are longer-lived than hermaphrodites, either on NGM plates (Gems and Riddle, 2000) or in liquid culture (McCulloch and Gems, 2003; McCulloch and Gems, 2007). Moreover, greater male longevity is a feature of many free-living nematodes (McCulloch and Gems, 2003). We attribute the difference between earlier observations and those in the present study to an unknown difference in culture conditions. We note that the mean lifespans of the controls are somewhat lower than usual in these trials.

**Table S8.** Effects of FUDR on lifespan when administered only after tumor development.

| Strain/<br>conditions | Number<br>of deaths/<br>censored | Mean<br>[median]<br>lifespan (days) | % change<br>vs.<br>N2 Carb | p vs. N2 Carb<br>(log rank) |
| --- | --- | --- | --- | --- |
| N2 Carb | [C] **85/5**<br>[1] 55/5<br>[2] 30/0 | **24.04 [24]**<br>25.42 [24]<br>21.50 [22] |  |  |
| N2<br>Carb + FUDR (D2) | [C] **120/0**<br>[1] 60/0<br>[2] 60/0 | **27.68 [28]**<br>29.05 [29.5]<br>26.30 [26.5] | **+15.14 [+16.67]**<br>+14.28 [+22.92]<br>+22.33 [+20.45] | **0.0003**<br>0.0053<br>0.0001 |
| N2<br>Carb + FUDR (D12) | [C] **108/12**<br>[1] 56/4<br>[2] 52/8 | **25.59 [26]**<br>25.84 [26]<br>25.33 [25] | **+6.45 [+8.33]**<br>+1.65 [+8.33]<br>+17.81 [+13.64] | **0.0284**<br>0.3655<br>0.0044 |
| N2<br>Carb + FUDR (D18) | [C] **110/10**<br>[1] 55/5<br>[2] 55/5 | **24.39 [24]**<br>22.42 [21]<br>26.36 [25] | **+1.46 [0]**<br>-11.80 [-12.50]<br>+22.60 [+13.64] | **0.3078**<br>0.1266<br>0.0002 |

**Table S9.** Effects of FUDR on lifespan in hermaphrodites and females of different free-living nematode species (*Caenorhabditis* species).

| Strain/<br>condition | Number of<br>deaths/<br>censored | Mean<br>[median]<br>lifespan<br>(days) | % change<br>vs.<br>control | p vs.<br>N2 control<br>(log rank) | % change<br>vs.<br>untreated | p vs.<br>untreated<br>(log rank) |
| --- | --- | --- | --- | --- | --- | --- |
| *C. tropicalis* (hermaphrodites) |  |  |  |  |  |  |
| Control | [C] 111/9<br>[1] 57/3<br>[2] 54/6 | 20.58 [21]<br>21.14 [21]<br>19.38 [21] |  |  |  |  |
| FUDR (D2) | [C] 88/32<br>[1] 39/21<br>[2] 49/11 | 19.95 [21]<br>18.67 [21]<br>20.98 [21] | -3.06 [0]<br>-11.68 [0]<br>+8.26 [0] | 0.5760<br>0.0267<br>0.4915 | -3.06 [0]<br>-11.68 [0]<br>+8.26 [0] | 0.5760<br>0.0267<br>0.4915 |
| Carb | [C] 106/14<br>[1] 55/5<br>[2] 51/9 | 20.25 [21]<br>21.87 [22]<br>18.49 [18] | -1.60 [0]<br>+3.45 [+4.76]<br>-4.59 [-14.29] | 0.4583<br>0.0746<br>0.2912 |  |  |
| Carb FUDR (D2) | [C] 98/22<br>[1] 48/12<br>[2] 50/10 | 24.46 [24]<br>25.73 [27]<br>23.24 [23] | +18.85 [+14.29]<br>+21.71 [+28.57]<br>+19.92 [+9.52] | <0.0001<br>0.0001<br>0.0089 | +20.79 [+14.29]<br>+17.65 [+22.73]<br>+25.69 [+27.78] | 0.0022<br>0.0959<br>0.0004 |
| *C. wallacei* (females) |  |  |  |  |  |  |
| Control | [C] 81/39<br>[1] 46/14<br>[2] 35/25 | 15.04 [14]<br>15.72 [15]<br>14.14 [13] |  |  |  |  |
| FUDR (D2) | [C] 75/15<br>[1] 28/2<br>[2] 47/13 | 17.13 [16]<br>18.07 [18]<br>16.57 [16] | +13.90 [+14.29]<br>+14.95 [+20.00]<br>+17.19 [+23.08] | 0.0127<br>0.0185<br>0.0731 | +13.90 [+14.29]<br>+14.95 [+20.00]<br>+17.19 [+23.08] | 0.0127<br>0.0185<br>0.0731 |
| Carb | [C] 89/31<br>[1] 46/14<br>[2] 43/17 | 20.28 [20]<br>22.63 [24]<br>17.77 [16] | +34.84 [+42.86]<br>+43.96 [+60.00]<br>+25.67 [+23.08] | <0.0001<br><0.0001<br>0.0127 |  |  |
| Carb FUDR (D2) | [C] 95/15<br>[1] 55/5<br>[2] 40/10 | 22.98 [24]<br>25.29 [27]<br>19.80 [16] | +52.79 [+71.43]<br>+60.88 [+80.00]<br>+40.03 [+23.08] | <0.0001<br><0.0001<br>0.0023 | +13.31 [+20.00]<br>+11.75 [+12.50]<br>+11.42 [0] | 0.0465<br>0.1424<br>0.3423 |

**Table S10.** Effects of FUDR on lifespan in hermaphrodites and females of different free-living nematode species (*Pristionchus* species).

| Strain/<br>condition | Number of<br>deaths/<br>censored | Mean<br>[median]<br>lifespan<br>(days) | % change<br>vs.<br>control | p vs.<br>N2 control<br>(log rank) | % change<br>vs.<br>untreated | p vs.<br>untreated<br>(log rank) |
| --- | --- | --- | --- | --- | --- | --- |
| *P. pacificus* (hermaphrodites) |  |  |  |  |  |  |
| control | [C] 122/3<br>[1] 30/0<br>[2] 34/1<br>[3] 58/2 | 25.61 [25]<br>26.03 [25]<br>25.06 [25]<br>25.72 [26] |  |  |  |  |
| FUDR (D2) | [C] 183/25<br>[1] 59/1<br>[2] 64/1<br>[3] 60/0 | 25.03 [25]<br>24.34 [25]<br>25.55 [25]<br>25.15 [25] | -2.26 [0]<br>-6.49 [0]<br>+1.96 [0]<br>-2.22 [-3.85] | 0.0324<br>0.0520<br>0.6329<br>0.4218 | -2.26 [0]<br>-6.49 [0]<br>+1.96 [0]<br>-2.22 [-3.85] | 0.0324<br>0.0520<br>0.6329<br>0.4218 |
| Carb | [C] 189/11<br>[1] 62/8<br>[2] 67/3<br>[3] 60/0 | 22.50 [22]<br>22.77 [23]<br>21.66 [21]<br>23.17 [22] | -12.14 [-12.00]<br>-12.52 [-8.00]<br>-13.57 [-16.00]<br>-9.91 [-15.38] | <0.0001<br>0.0023<br><0.0001<br>0.0004 |  |  |
| Carb FUDR (D2) | [C] 197/3<br>[1] 68/2<br>[2] 69/1<br>[3] 60/0 | 24.10 [24]<br>23.65 [25]<br>24.10 [25]<br>24.62 [24] | -5.90 [-4.00]<br>-9.14 [0]<br>-3.83 [0]<br>-4.28 [-7.69] | <0.0001<br>0.0038<br>0.0286<br>0.0652 | +7.11 [+9.09]<br>+3.86 [+8.70]<br>+11.27 [+19.05]<br>+6.26 [+9.09] | 0.0001<br>0.2894<br><0.0001<br>0.0366 |
| *P. exspectatus* (females) |  |  |  |  |  |  |
| control | [C] 121/24<br>[1] 22/8<br>[2] 51/9<br>[3] 48/7 | 36.59 [36]<br>35.18 [37]<br>39.00 [39]<br>34.67 [36] |  |  |  |  |
| FUDR (D2) | [C] 155/15<br>[1] 39/11<br>[2] 59/1<br>[3] 57/3 | 31.69 [31]<br>31.67 [30]<br>33.14 [32]<br>30.21 [31] | -13.39 [-13.89]<br>-9.98 [-18.92]<br>-15.03 [-17.95]<br>-12.86 [-13.89] | <0.0001<br>0.0179<br><0.0001<br>0.0011 | -13.39 [-13.89]<br>-9.98 [-18.92]<br>-15.03 [-17.95]<br>-12.86 [-13.89] | <0.0001<br>0.0179<br><0.0001<br>0.0011 |
| Carb | [C] 127/28<br>[1] 40/10<br>[2] 49/11<br>[3] 38/7 | 36.52 [37]<br>37.18 [37]<br>38.88 [39]<br>32.97 [33] | -0.19 [+2.78]<br>+5.69 [0]<br>-0.31 [0]<br>-4.90 [-8.33] | 0.7435<br>0.4291<br>0.6199<br>0.4102 |  |  |
| Carb FUDR (D2) | [C] 150/15<br>[1] 50/10<br>[2] 58/2<br>[3] 42/3 | 30.80 [31]<br>30.32 [32]<br>31.38 [31]<br>30.57 [31] | -15.82 [-13.89]<br>-13.81 [-13.51]<br>-19.54 [-20.51]<br>-11.83 [-13.89] | <0.0001<br>0.0047<br><0.0001<br>0.0108 | -15.66 [-16.22]<br>-18.45 [-13.51]<br>-19.29 [-20.51]<br>-7.28 [-6.06] | <0.0001<br><0.0001<br><0.0001<br>0.0765 |

**Table S11.** Effects of *vit-5,-6* RNAi, Kan and 50 μM FUDR on senescent pathologies (statistical comparisons).

| Strain/<br>conditions | Time | Pharyngeal deterioration |  |  | Distal gonad degeneration |  |  | Uterine tumor |  |  | Intestinal atrophy |  |  | Yolk pool accumulation |  |  |
| --- | --- | --- | --- | --- | --- | --- | --- | --- | --- | --- | --- | --- | --- | --- | --- | --- |
|  |  | *p* vs.<br>non<br>FUDR | *p* vs.<br>L4440<br>treatment | *p* vs.<br>non Kan | *p* vs.<br>non<br>FUDR | *p* vs.<br>L4440<br>treatment | *p* vs.<br>non Kan | *p* vs.<br>non<br>FUDR | *p* vs.<br>L4440<br>treatment | *p* vs.<br>non Kan | *p* vs.<br>non<br>FUDR | *p* vs.<br>L4440<br>treatment | *p* vs.<br>non Kan | *p* vs.<br>non<br>FUDR | *p* vs.<br>L4440<br>treatment | *p* vs.<br>non Kan |
| L4440 | Day 1 |  |  |  |  |  |  |  |  |  |  |  |  |  |  |  |
|  | Day 7 |  |  |  |  |  |  |  |  |  |  |  |  |  |  |  |
|  | Day14 |  |  |  |  |  |  |  |  |  |  |  |  |  |  |  |
| *vit-5,-6* RNAi | Day 1 |  | >0.99 |  |  | 0.99 |  |  | >0.99 |  |  | 0.81 |  |  | >0.99 |  |
|  | Day 7 |  | 0.99 |  |  | 0.32 |  |  | **<0.0001** |  |  | **0.0022** |  |  | 0.106 |  |
|  | Day14 |  | 0.47 |  |  | 0.42 |  |  | **<0.0001** |  |  | **0.0010** |  |  | **0.0023** |  |
| L4440 FUDR | Day 1 | >0.99 |  |  | 0.37 |  |  | >0.99 |  |  | **0.028** |  |  | 0.99 |  |  |
|  | Day 7 | >0.99 |  |  | >0.99 |  |  | **<0.0001** |  |  | 0.83 |  |  | **0.033** |  |  |
|  | Day14 | **0.0023** |  |  | **<0.0001** |  |  | 0.72 |  |  | 0.35 |  |  | **0.0069** |  |  |
| *vit-5,-6* RNAi<br>FUDR | Day 1 | >0.99 | >0.99 |  | 0.81 | 0.97 |  | >0.99 | >0.99 |  | **0.0089** | 0.82 |  | 0.88 | 0.91 |  |
|  | Day 7 | >0.99 | 0.99 |  | 0.54 | 0.95 |  | **<0.0001** | **0.032** |  | **0.0003** | 0.83 |  | *0.099* | 0.59 |  |
|  | Day14 | 0.44 | 0.93 |  | **0.0002** | 0.25 |  | 0.98 | 0.47 |  | 0.98 | *0.068* |  | **0.0008** | **0.0028** |  |
| L4440 Kan | Day 1 |  |  | >0.99 |  |  | 0.23 |  |  | >0.99 |  |  | 0.98 |  |  | 0.76 |
|  | Day 7 |  |  | 0.93 |  |  | 0.84 |  |  | 0.84 |  |  | 0.95 |  |  | >0.99 |
|  | Day14 |  |  | 0.91 |  |  | 0.52 |  |  | 0.26 |  |  | 0.99 |  |  | 0.81 |
| *vit-5,-6* Kan | Day 1 |  | >0.99 | >0.99 |  | 0.48 | 0.99 |  | >0.99 | >0.99 |  | 0.4831 | 0.18 |  | 0.95 | 0.92 |
|  | Day 7 |  | 0.99 | 0.99 |  | 0.18 | 0.98 |  | **0.0001** | 0.95 |  | 0.8623 | **0.0008** |  | **0.012** | 0.88 |
|  | Day14 |  | 0.27 | 0.75 |  | 0.40 | 0.58 |  | 0.21 | 0.90 |  | **0.0381** | 0.65 |  | 0.308 | **0.015** |
| L4440<br>Kan + FUDR | Day 1 | >0.99 |  | >0.99 | 0.90 |  | 0.67 | >0.99 |  | >0.99 | **0.021** |  | 0.89 | 0.92 |  | 0.91 |
|  | Day 7 | 0.90 |  | >0.99 | 0.97 |  | 0.65 | **<0.0001** |  | 0.43 | 0.85 |  | 0.89 | **0.0001** |  | 0.86 |
|  | Day 14 | **<0.0001** |  | 0.32 | **0.0007** |  | 0.64 | 0.80 |  | 0.94 | 0.23 |  | >0.99 | **<0.0001** |  | >0.99 |
| *vit-5,-6* RNAi<br>Kan + FUDR | Day 1 | >0.99 | >0.99 | >0.99 | 0.52 | 0.77 | 0.91 | >0.99 | >0.99 | >0.99 | 0.81 | 0.90 | 0.97 | 0.93 | 0.89 | 0.85 |
|  | Day 7 | 0.98 | >0.99 | 0.99 | 0.87 | 0.27 | 0.99 | **<0.0001** | 0.38 | 0.99 | 0.49 | 0.44 | 0.49 | 0.94 | 0.99 | 0.80 |
|  | Day 14 | 0.30 | 0.91 | 0.52 | **0.016** | 0.93 | >0.99 | 0.39 | **0.026** | 0.85 | 0.55 | *0.062* | >0.99 | **<0.0001** | **0.0144** | 0.79 |

Bold, $p < 0.05$ ; italics, $0.1 > p > 0.05$ .

**Table S12.** Combined effects of blocking infection, tumor development and vitellogenesis (20°C)

| Strain/<br>conditions | Number<br>of deaths/<br>censored | Mean<br>[Median]<br>lifespan (days) | % change<br>vs. L4440 | *p* vs. L4440<br>(log rank) | % change<br>vs.<br>L4440 treatment | *p* vs.<br>L4440 treatment<br>(log rank) | *p* vs.<br>L4440 treatment<br>(CPH test) |
| --- | --- | --- | --- | --- | --- | --- | --- |
| L4440 | [C] 182/16<br>[1] 57/5<br>[2] 56/7<br>[3] 69/4 | 17.09 [17]<br>17.53 [17]<br>17.25 [17]<br>16.59 [17] |  |  |  |  |  |
| *vit-5,-6* RNAi | [C] 175/22<br>[1] 57/7<br>[2] 59/9<br>[3] 59/6 | 20.58 [21]<br>20.02 [21]<br>20.81 [21]<br>20.88 [21] | +20.42 [+23.53]<br>+14.20 [+23.53]<br>+20.64 [+23.53]<br>+25.86 [+23.53] | <0.0001<br>0.0096<br>0.0007<br>0.0001 |  |  |  |
| L4440 FUDR | [C] 192/7<br>[1] 62/0<br>[2] 63/1<br>[3] 67/6 | 17.14 [15]<br>17.65 [15]<br>17.83 [18]<br>16.03 [15] | +0.29 [-11.76]<br>+0.68 [-11.76]<br>+3.36 [+5.88]<br>-3.38 [-11.76] | 0.7062<br>0.9728<br>0.8844<br>0.5164 |  |  |  |
| *vit-5,-6* RNAi<br>FUDR | [C] 177/5<br>[1] 65/0<br>[2] 53/1<br>[3] 59/4 | 21.97 [23]<br>22.43 [23]<br>22.82 [23]<br>20.69 [21] | +28.55 [+35.29]<br>+27.95 [+35.29]<br>+32.29 [+35.29]<br>+24.71 [+23.53] | <0.0001<br><0.0001<br><0.0001<br><0.0001 | +28.18 [+53.33]<br>+27.08 [+53.33]<br>+27.99 [+27.78]<br>+29.07 [+40.00] | <0.0001<br><0.0001<br><0.0001<br><0.0001 | 0.0133<br>0.0363<br>0.1656<br>0.8582 |
| L4440 Kan | [C] 189/22<br>[1] 63/8<br>[2] 61/9<br>[3] 65/5 | 22.08 [21]<br>23.84 [25]<br>22.28 [21]<br>20.20 [19] | +29.20 [+23.53]<br>+36.00 [+47.06]<br>+29.16 [+23.53]<br>+21.76 [+11.76] | <0.0001<br><0.0001<br><0.0001<br>0.0006 |  |  |  |
| *vit-5,-6* RNAi<br>Kan | [C] 184/17<br>[1] 57/5<br>[2] 63/5<br>[3] 64/7 | 23.26 [23]<br>23.11 [23]<br>22.60 [21]<br>24.03 [23] | +36.10 [+35.29]<br>+31.83 [+35.29]<br>+31.01 [+23.53]<br>+44.85 [+35.29] | <0.0001<br><0.0001<br><0.0001<br><0.0001 | +5.34 [+9.52]<br>-3.06 [-8.00]<br>+1.44 [0]<br>+18.96 [+21.05] | 0.0190<br>0.9192<br>0.3177<br>0.0008 | 0.0659<br>0.0974<br>0.2023<br>0.9379 |
| L4440<br>Kan + FUDR | [C] 182/12<br>[1] 60/3<br>[2] 62/3<br>[3] 60/6 | 23.43 [23]<br>23.82 [25]<br>23.15 [23]<br>23.35 [23] | +37.10 [+35.29]<br>+35.88 [+47.06]<br>+34.20 [+35.29]<br>+40.75 [+35.29] | <0.0001<br><0.0001<br><0.0001<br><0.0001 |  |  |  |
| *vit-5,-6* RNAi<br>Kan + FUDR | [C] 191/13<br>[1] 59/5<br>[2] 71/2<br>[3] 61/6 | 24.69 [25]<br>23.88 [25]<br>25.35 [27]<br>24.74 [25] | +44.47 [+47.06]<br>+36.22 [+47.06]<br>+43.42 [+58.82]<br>+49.13 [+47.06] | <0.0001<br><0.0001<br><0.0001<br><0.0001 | +5.38 [+8.70]<br>+0.25 [+0]<br>+9.50 [+17.39]<br>+5.95 [+8.70] | 0.0140<br>0.5270<br>0.0330<br>0.2210 | 0.0241<br>0.5851<br>0.9309<br>0.0489 |

**Table S13.** Prevention of *E. coli* infection causes *ced-3* to shorten lifespan.

| Strain/<br>conditions | Number<br>of deaths/<br>censored | Mean<br>[median]<br>lifespan<br>(days) | % change<br>vs.<br>N2 control | p vs. N2<br>(log rank) | % change<br>vs.<br>N2 Carb | p vs.<br>N2 Carb<br>(log rank) |
| --- | --- | --- | --- | --- | --- | --- |
| N2 control | [C] 126/26<br>[1] 46/13<br>[2] 56/4<br>[3] 23/9 | 17.46 [17]<br>16.74 [17]<br>18.62 [17]<br>16.04 [16] |  |  |  |  |
| *ced-3(n717)* | [C] 132/20<br>[1] 53/7<br>[2] 56/4<br>[3] 23/9 | 18.47 [18]<br>18.52 [17]<br>18.40 [20]<br>18.60 [18] | +5.78 [+5.88]<br>+10.63 [0]<br>-1.18 [+17.65]<br>+15.96 [+12.50] | 0.1226<br>0.0415<br>0.7763<br>0.1401 |  |  |
| *ced-3(n1286)* | [C] 110/43<br>[1] 48/12<br>[2] 41/19<br>[3] 21/12 | 17.97 [18]<br>17.41 [17]<br>18.79 [20]<br>17.62 [16] | +2.92 [+5.88]<br>+4.00 [0]<br>+0.91 [+17.65]<br>+9.85 [0] | 0.3327<br>0.2474<br>0.7951<br>0.2799 |  |  |
| *ced-3(n2454)* | [C] 103/22<br>[1] 55/5<br>[2] 25/5<br>[3] 23/8 | 18.04 [19]<br>17.75 [19]<br>18.02 [20]<br>18.74 [18] | +3.32 [+11.76]<br>+6.03 [+11.76]<br>-3.22 [+17.65]<br>+16.83 [+12.50] | 0.7851<br>0.3895<br>0.2628<br>0.1975 |  |  |
| N2 Carb | [C] 167/24<br>[1] 52/8<br>[2] 57/3<br>[3] 58/13 | 26.28 [27]<br>25.83<br>[26.5]<br>27.68 [27]<br>25.31 [25] | +50.52 [+58.82]<br>+54.30 [+55.88]<br>+48.66 [+58.82]<br>+57.79 [+56.25] | <0.0001<br><0.0001<br><0.0001<br><0.0001 |  |  |
| *ced-3(n717)* Carb | [C] 140/41<br>[1] 45/15<br>[2] 54/6<br>[3] 41/20 | 23.42 [22]<br>23.31 [21]<br>23.52 [22]<br>23.40 [23] | +34.14 [+29.41]<br>+39.25 [+23.53]<br>+26.32 [+29.41]<br>+45.89 [+43.75] | <0.0001<br><0.0001<br><0.0001<br><0.0001 | -10.88 [-18.52]<br>-9.76 [-20.75]<br>-15.03 [-18.52]<br>-7.55 [-8.00] | <0.0001<br>0.0284<br>0.0002<br>0.0347 |
| *ced-3(n1286)*<br>Carb | [C] 103/47<br>[1] 39/21<br>[2] 41/19<br>[3] 23/7 | 23.30 [23]<br>22.96 [21]<br>23.97 [24]<br>22.91 [23] | +33.45 [+35.29]<br>+37.16 [+23.53]<br>+28.73 [+41.18]<br>+42.83 [+43.75] | <0.0001<br><0.0001<br><0.0001<br>0.0002 | -11.34 [-14.81]<br>-11.11 [-20.75]<br>-13.40 [-11.11]<br>-9.48 [-8.00] | <0.0001<br>0.0249<br>0.0018<br>0.0361 |
| *ced-3(n2454)*<br>Carb | [C] 127/64<br>[1] 47/13<br>[2] 50/10<br>[3] 30/41 | 22.04 [22]<br>22.15 [21]<br>22.19 [22]<br>21.64 [21] | +26.23 [+29.41]<br>+32.32 [+23.53]<br>+19.17 [+29.41]<br>+34.91 [+31.25] | <0.0001<br><0.0001<br>0.0189<br>0.0007 | -16.13 [-18.52]<br>-14.25 [-20.75]<br>-19.83 [-18.52]<br>-14.50 [-16.00] | <0.0001<br>0.0043<br><0.0001<br>0.0024 |

**Table S14.**
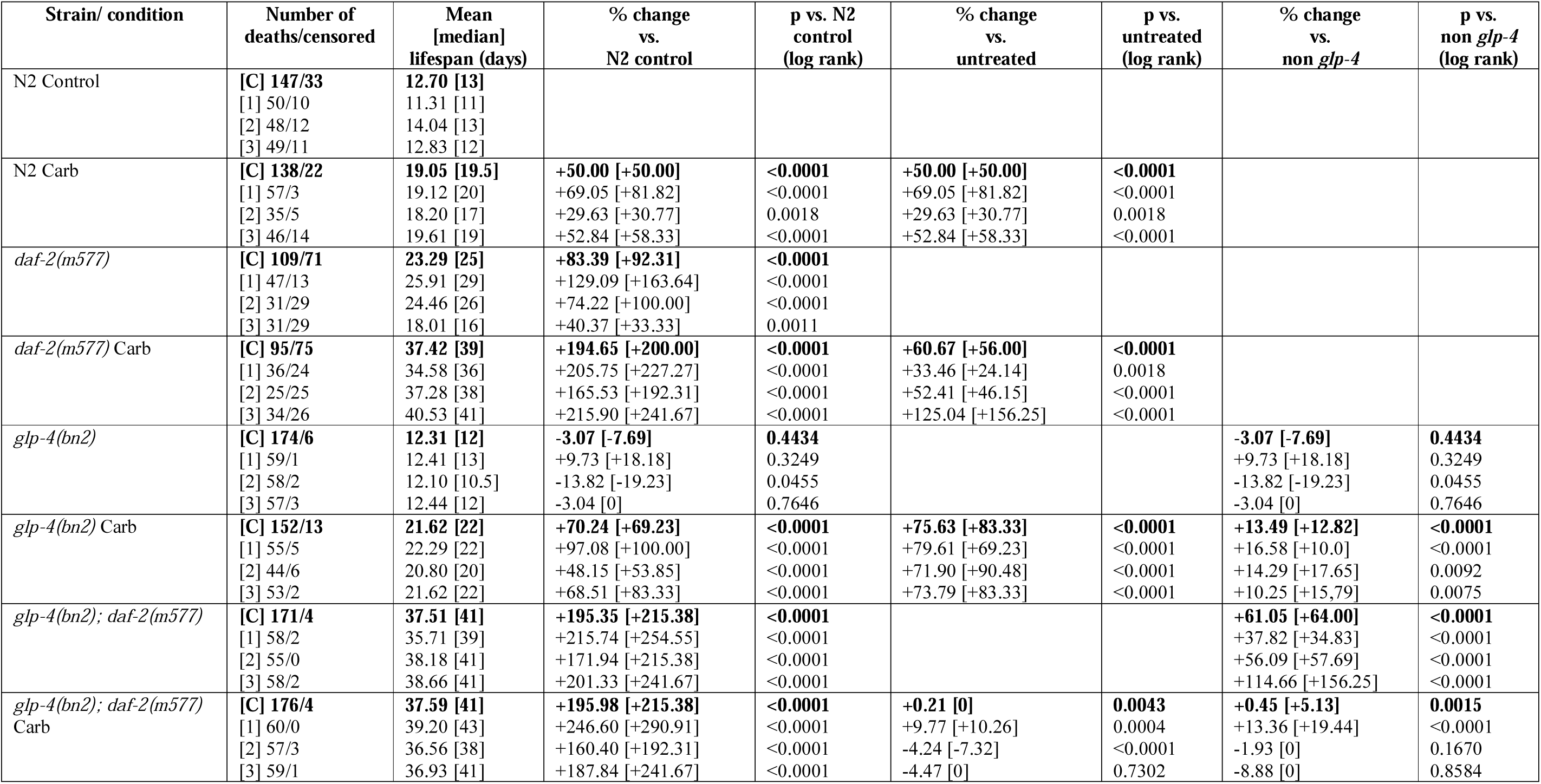
Carb suppresses enhancement of *daf-2* longevity by *glp-4(bn2)* (25°C from L4 stage)

| Strain/ condition | Number of deaths/censored | Mean [median] lifespan (days) | % change vs. N2 control | p vs. N2 control (log rank) | % change vs. untreated | p vs. untreated (log rank) | % change vs. non *glp-4* | p vs. non *glp-4* (log rank) |
| --- | --- | --- | --- | --- | --- | --- | --- | --- |
| N2 Control | [C] 147/33<br>[1] 50/10<br>[2] 48/12<br>[3] 49/11 | 12.70 [13]<br>11.31 [11]<br>14.04 [13]<br>12.83 [12] |  |  |  |  |  |  |
| N2 Carb | [C] 138/22<br>[1] 57/3<br>[2] 35/5<br>[3] 46/14 | 19.05 [19.5]<br>19.12 [20]<br>18.20 [17]<br>19.61 [19] | +50.00 [+50.00]<br>+69.05 [+81.82]<br>+29.63 [+30.77]<br>+52.84 [+58.33] | <0.0001<br><0.0001<br>0.0018<br><0.0001 | +50.00 [+50.00]<br>+69.05 [+81.82]<br>+29.63 [+30.77]<br>+52.84 [+58.33] | <0.0001<br><0.0001<br>0.0018<br><0.0001 |  |  |
| *daf-2(m577)* | [C] 109/71<br>[1] 47/13<br>[2] 31/29<br>[3] 31/29 | 23.29 [25]<br>25.91 [29]<br>24.46 [26]<br>18.01 [16] | +83.39 [+92.31]<br>+129.09 [+163.64]<br>+74.22 [+100.00]<br>+40.37 [+33.33] | <0.0001<br><0.0001<br><0.0001<br>0.0011 |  |  |  |  |
| *daf-2(m577)* Carb | [C] 95/75<br>[1] 36/24<br>[2] 25/25<br>[3] 34/26 | 37.42 [39]<br>34.58 [36]<br>37.28 [38]<br>40.53 [41] | +194.65 [+200.00]<br>+205.75 [+227.27]<br>+165.53 [+192.31]<br>+215.90 [+241.67] | <0.0001<br><0.0001<br><0.0001<br><0.0001 | +60.67 [+56.00]<br>+33.46 [+24.14]<br>+52.41 [+46.15]<br>+125.04 [+156.25] | <0.0001<br>0.0018<br><0.0001<br><0.0001 |  |  |
| *glp-4(bn2)* | [C] 174/6<br>[1] 59/1<br>[2] 58/2<br>[3] 57/3 | 12.31 [12]<br>12.41 [13]<br>12.10 [10.5]<br>12.44 [12] | -3.07 [-7.69]<br>+9.73 [+18.18]<br>-13.82 [-19.23]<br>-3.04 [0] | 0.4434<br>0.3249<br>0.0455<br>0.7646 |  |  | -3.07 [-7.69]<br>+9.73 [+18.18]<br>-13.82 [-19.23]<br>-3.04 [0] | 0.4434<br>0.3249<br>0.0455<br>0.7646 |
| *glp-4(bn2)* Carb | [C] 152/13<br>[1] 55/5<br>[2] 44/6<br>[3] 53/2 | 21.62 [22]<br>22.29 [22]<br>20.80 [20]<br>21.62 [22] | +70.24 [+69.23]<br>+97.08 [+100.00]<br>+48.15 [+53.85]<br>+68.51 [+83.33] | <0.0001<br><0.0001<br><0.0001<br><0.0001 | +75.63 [+83.33]<br>+79.61 [+69.23]<br>+71.90 [+90.48]<br>+73.79 [+83.33] | <0.0001<br><0.0001<br><0.0001<br><0.0001 | +13.49 [+12.82]<br>+16.58 [+10.0]<br>+14.29 [+17.65]<br>+10.25 [+15.79] | <0.0001<br><0.0001<br>0.0092<br>0.0075 |
| *glp-4(bn2); daf-2(m577)* | [C] 171/4<br>[1] 58/2<br>[2] 55/0<br>[3] 58/2 | 37.51 [41]<br>35.71 [39]<br>38.18 [41]<br>38.66 [41] | +195.35 [+215.38]<br>+215.74 [+254.55]<br>+171.94 [+215.38]<br>+201.33 [+241.67] | <0.0001<br><0.0001<br><0.0001<br><0.0001 |  |  | +61.05 [+64.00]<br>+37.82 [+34.83]<br>+56.09 [+57.69]<br>+114.66 [+156.25] | <0.0001<br><0.0001<br><0.0001<br><0.0001 |
| *glp-4(bn2); daf-2(m577)* Carb | [C] 176/4<br>[1] 60/0<br>[2] 57/3<br>[3] 59/1 | 37.59 [41]<br>39.20 [43]<br>36.56 [38]<br>36.93 [41] | +195.98 [+215.38]<br>+246.60 [+290.91]<br>+160.40 [+192.31]<br>+187.84 [+241.67] | <0.0001<br><0.0001<br><0.0001<br><0.0001 | +0.21 [0]<br>+9.77 [+10.26]<br>-4.24 [-7.32]<br>-4.47 [0] | 0.0043<br>0.0004<br><0.0001<br>0.7302 | +0.45 [+5.13]<br>+13.36 [+19.44]<br>-1.93 [0]<br>-8.88 [0] | 0.0015<br><0.0001<br>0.1670<br>0.8584 |

**Table S15.**
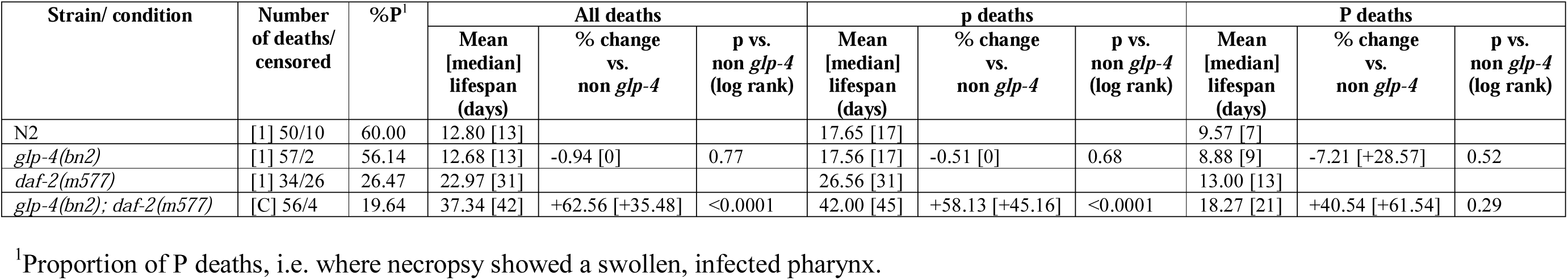
Mortality deconvolution analysis of effects of *glp-4(bn2)* on lifespan in *daf-2(m577)* (no Carb)

